# Ambient temperature structural studies of Penicillin-binding Protein 2a of *Methicillin-Resistant Staphylococcus aureus* with XFELs and synchrotrons

**DOI:** 10.1101/2025.02.22.637770

**Authors:** Alice Grieco, Isabel Quereda-Moraleda, Sabine Botha, Katerina Dörner, Christina Schmidt, Huijong Han, Mohammad Vakili, Joachim Schulz, Faisal Koua, Raphael de Wijn, Konstantin Kharitonov, Adam Round, Johan Bielecki, Christopher Kupitz, Stella Lisova, Pamela Schleissner, Ray Sierra, Valerio Mariani, Shibom Basu, Julien Orlans, Samuel Rose, Daniele de Sanctis, Leland B. Gee, Richard Bean, Shahriar Mobashery, Mayland Chang, Juan A. Hermoso, Jose Manuel Martin-Garcia

## Abstract

Penicillin-binding protein 2a (PBP2a) is a transpeptidase responsible for the β-lactam resistance in methicillin-resistant *Staphylococcus aureus* (MRSA), posing significant challenges to antibiotic therapy. PBP2a’s unique structural features, including its highly flexible active site and allosteric regulation, enable it to maintain catalytic activity even in the presence of β-lactam antibiotics. Despite extensive characterization using cryogenic crystallography, key questions remain about its dynamic properties and conformational changes under near-physiological conditions. Room-temperature crystallography methods, particularly serial femtosecond X-ray crystallography (SFX) at XFELs, provide a powerful approach to capture these dynamics. Unlike cryogenic conditions that can constrain protein flexibility, SFX enables the study of conformational variability and interaction networks closer to physiological states. Here, we present the first room-temperature structures of PBP2a obtained using SFX to uncover insights into the enzyme’s flexibility, allosteric communication, and catalytic mechanisms. These findings are built upon optimized large-scale production and crystallization protocols for PBP2a, ensuring high-quality microcrystals suitable for SFX data collection. The room-temperature structures reveal novel interaction patterns, including unique salt bridge networks and dynamic structural elements absent in cryogenic studies. Furthermore, comparative analyses highlight how environmental conditions influence the conformational states of PBP2a, providing new perspectives on its resistance mechanisms. By integrating structural data from EuXFEL and LCLS, this study not only enhances our understanding of PBP2a’s functional dynamics but also underscores the value of room-temperature crystallography in studying antibiotic resistance. These insights could guide the design of next-generation β-lactam antibiotics capable of overcoming PBP2a-mediated resistance.

## 1. Introduction

*Staphylococcus aureus*, a Gram-positive bacterium, remains a significant public health threat due to its resistance to major antibacterial therapies. *Methicillin-resistant S. aureus* (MRSA) is particularly problematic, ranking among the most common global causes of infective endocarditis, as well as skin and respiratory infections (Abdel-Moein & Zaher, 2019; Rahman *et al*., 2018). Many countries have implemented stringent antibiotic use policies to preserve the efficacy of existing treatments (Stryjewski & Corey, 2014). Despite these efforts, reports of MRSA strains with reduced sensitivity or complete resistance to current antibiotics, such as daptomycin, ceftaroline, vancomycin, amoxicillin, linezolid, and cefotaxime, are on the rise (Stryjewski & Corey, 2014; Malik & Bhattacharyya, 2019; Verma *et al*., 2022; Larsson & Flach, 2022). Bacterial survival relies heavily on the integrity of their cell walls, primarily composed of peptidoglycan (Josephine *et al*., 2006; Zapun *et al*., 2008). Peptidoglycan is made up of repeating units of a disaccharide, N-acetyl glucosamine (NAG) and N-acetylmuramic acid (NAM), with peptide stems attached to the NAM units. These stems crosslink with those of adjacent peptidoglycan units to form a mature cell wall (Sauvage *et al*., 2008; Dirk-Jan & G., 2005). In *S. aureus*, the peptide stem consists of a five-chain peptide: L-Ala-γ-D-Glu-L-Lys(Gly)5-D-Ala-D-Ala, where the lysine side chain is linked to pentaglycine during cell wall crosslinking (Sauvage *et al*., 2008). The transglycosylase activity, responsible for synthesizing the peptidoglycan backbone, and transpeptidase activity, which facilitates peptidoglycan crosslinking, are both carried out by penicillin-binding proteins (PBPs) (Sauvage *et al*., 2008). Given their essential role in cell wall synthesis and bacterial survival, PBPs serve as excellent targets for antibiotics, particularly β-lactams. Therefore, in addition to promoting safe antibiotic practices, developing new therapeutics with novel mechanisms of action is a top priority for antimicrobial drug development research groups to combat the escalating issue of antibiotic resistance.

Penicillin-binding protein 2a (PBP2a) is a key enzyme in MRSA, conferring resistance to β-lactam antibiotics, which are commonly used to treat bacterial infections. PBP2a is an altered form of the PBPs found in non-resistant *S. aureus* strains, which play a crucial role in bacterial cell wall synthesis by catalyzing the cross-linking of peptidoglycan strands (Yao *et al*., 2012), an essential process for maintaining cell integrity (Kim *et al*., 2012). Unlike these typical PBPs, which have a high affinity for β-lactam antibiotics and are readily inhibited by them, PBP2a has significantly reduced affinity for β-lactam antibiotics, allowing MRSA to continue synthesizing its cell wall in the presence of these drugs, thereby surviving and proliferating despite antibiotic treatment (Fuda *et al*., 2004). Structural information on PBP2a is crucial for fully understanding its enzymatic process. In this regard, previous crystallographic structures at cryogenic temperature have reported the presence of an allosteric binding domain 60 Å distant from the transpeptidase active site. When this allosteric site is occupied, multiresidue conformational changes culminates in the opening of the active site to permit substrate entry and transpeptidation (Otero *et al*., 2013). X-ray crystallography experiments have identified two conformational states (open and closed) for the residues interconnecting the allosteric and active sites. Importantly, ligand-bound structures of PBP2a have been reported, providing valuable insights into how binding at the allosteric site influences active-site accessibility. These structures support the idea that ligand binding can induce structural rearrangements, which may be crucial for designing novel inhibitors that target PBP2a’s unique regulatory mechanism. However, kinetic information characterizing the dynamic transitions between these conformations remains unknown (Otero *et al*., 2013).

Despite its success in determining 3D structures of macromolecules, standard single-crystal macromolecular X-ray crystallography typically requires large, well-ordered crystals and operates under nonphysiologically, cryogenic conditions, which indeed can obscure biologically relevant conformational states (Fraser *et al*., 2011). In contrast, room temperature crystallography preserves the native physiological conditions of biological macromolecules, enabling our ability to capture dynamic processes and conformational flexibility, and providing unprecedented structural insights into their true functional states (Keedy *et al*., 2015). In this regard, time-resolved studies using serial femtosecond X-ray crystallography (TR-SFX) at X-ray free-electron lasers (XFELs) have successfully investigated enzyme-catalyzed reactions at atomic resolution. In these studies, hundreds of thousands of diffraction snapshots are collected from microcrystals at room temperature (Chapman *et al*., 2011). These crystals interact with ultrabright, ultrashort X-ray pulses of 10–40 fs duration. Despite the intense pulse intensity, the pulses are short enough to collect diffraction images before secondary X-ray radiation damage occurs. This enables the determination of radiation-free macromolecular structures at ambient temperature. Using mix-and-inject TR-SFX technology, biological reactions can be triggered "on the fly" to study protein mechanisms *in real time*. This method has been applied successfully to study the enzymatic function of β-lactamase C (BlaC) from *Mycobacterium tuberculosis* (Kupitz *et al*., 2017; Olmos *et al*., 2018; Pandey *et al*., 2021) and to perform dynamic studies of an RNA riboswitch (Stagno *et al*., 2017).

In the study presented, we propose the structural analysis of the PBP2a from MRSA both at synchrotrons and XFELs (Llarrull *et al*., 2009). The ultimate goal of our research will be to solve the crystallographic structures of the intermediates involved in the PBP2a reaction mechanism. To this end and in preparation for future TR-SFX studies, we present here, to the best of our knowledge, the first room-temperature structures of PBP2a obtained using SFX, comparing them to cryogenic structures to uncover insights into the enzyme’s flexibility, allosteric communication, and catalytic mechanisms. These findings are built upon optimized large-scale production and crystallization protocols for PBP2a, ensuring high-quality microcrystals suitable for SFX data collection. The room-temperature structures reveal novel interaction patterns, including unique salt bridge networks and dynamic structural elements absent in cryogenic studies. Furthermore, comparative analyses highlight how environmental conditions influence the conformational states of PBP2a, providing new perspectives on its resistance mechanisms. By integrating structural data from EuXFEL and LCLS, this study not only enhances our understanding of PBP2a’s functional dynamics but also underscores the value of room-temperature crystallography in studying antibiotic resistance. These insights could guide the design of next-generation β-lactam antibiotics capable of overcoming PBP2a-mediated resistance.

## 2. Materials and methods

### 2.1 Materials

*Escherichia coli* BL21 (DE3) competent cells were purchased from Novagen. Yeast extract and tryptone were purchased from Condalab (Madrid, Spain). EDTA-free protease inhibitor cocktail, isopropyl β-d-1-thiogalactopyranoside (IPTG), Kanamycin, ammonium sulfate, sodium chloride and Tris-HCl were purchased from Merck (Madrid, Spain). HEPES, Polyethylene glycol (PEG) 3350 and 1500 were purchased from Sigma-Aldrich. Macro Prep Hight S Support column was purchased from Cytiva and HiLoad 16/600 Superdex 200 from GE Healthcare.

### 2.2. Expression and purification of PBP2a

*Escherichia coli* BL21(DE3) cells were transformed with pET24a(+)-mecA plasmid containing the cDNA of the penicillin binding protein 2a (PBP2a) of *Staphylococcus aureus* ATCC706986 without the first 22 residues at the N-terminal corresponding to the transmembrane portion of the protein. Transformed cells were grown at 37 °C in Luria Bertani (LB) medium containing kanamycin (50 µg/ mL) until exponential phase reached an OD_600_ equal to 0.6, followed by induction with 0.5 mM isopropyl-β-D-thiogalactopyranoside (IPTG). Induced cells were further incubated overnight at 25 °C, harvested by centrifugation at 6,000 rpm at 4 °C for 15 min, resuspended in 10 mL of freshly prepared buffer A (20 mM Tris-HCl pH 8, 5 μL of DNase in water (1mg/mL), 1 tablet of EDTA-free inhibitors, 5 mM MgCl2), flash frozen in liquid N_2_, and stored at −80 °C. The frozen cells were thawed and lysed by sonication using 3 cycles of 2 min each, alternating 2 s ON and 2 s OFF with 2 min rest on ice. The lysate was cleared by centrifugation at 30,000 rpm at 4 °C for 40 min. The supernatant containing PBP2a was transferred to a glass beaker containing a stir bar and placed on a magnetic stirrer. While the sample was stirring, ammonium sulfate powder was slowly added to bring the final concentration to 65% saturation. Once the ammonium sulfate was added, the sample was left in continuous agitation for one hour at 4°C. Then, the sample was spun down at 30,000 rpm at 4 °C for 40 min to remove the precipitate. The supernatant was removed and transferred to a dialysis bag and extensively dialyzed (3 exchanges of 5L each) against buffer B (25 mM HEPES pH 7, 200 mM NaCl). The following day, the sample was removed from the dialysis and loaded onto an Ion exchange chromatography Macro Prep Hight S Support column (CYTIVA, Marlborough, USA), which was previously equilibrated with 20 column volumes (CV) of buffer B. After collecting the flow through, the column was washed with 4 CV of buffer B and eluted with 4 CV of elution buffer C (25 mM HEPES pH 7, 1 M NaCl). The eluted protein was further purified by size-exclusion chromatography (SEC) using a HiLoad 16/600 Superdex 200 prep grade (GE Healthcare) column using buffer D (25 mM HEPES pH 7, 100 mM NaCl) as elution buffer. Pure protein was concentrated to a final concentration of 30 mg/mL using concentrators from Millipore with a MW cut-off of 30 kDa, flash frozen in liquid N_2_ and stored at −80 °C. The purity and integrity of the protein were checked by SDS-PAGE.

### 2.3. Analytical ultracentrifugation

Experiments were conducted using an Optima XL-A analytical ultracentrifuge (Beckman-Coulter Inc.) equipped with UV-VIS absorbance detection system. An An-50Ti rotor and 12 mm optical path epon-charcoal standard double-sector centerpieces were used. Samples of PBP2a were analyzed in four different buffers: 1) 20 mM HEPES, pH 7.0, 150 mM NaCl; 2) 20 mM HEPES, pH 7.0, 150 mM NaCl, 20 mM CdCl2; 3) 20 mM HEPES, pH 7.0, 150 mM NaCl, 20 mM CdCl2, 20 mM EDTA; 4) 20 mM HEPES, pH 7.0, 500 mM NaCl. For buffer 1, assays were performed at three protein concentrations: 0.5, 1.0, and 1.5 mg/mL. For buffers 2, 3, and 4, ultracentrifugation experiments were conducted at a single protein concentration of 1.0 mg/mL. All samples were centrifuged at 48,000 rpm at 20°C. Sedimentation was monitored by absorbance at 280 nm.

Differential sedimentation coefficient distributions were determined by least-squares boundary modeling of sedimentation velocity data using the continuous distribution c(s) Lamm equation model implemented in the SEDFIT software (Schuck, 2000). Experimental sedimentation coefficient values (s) were corrected to standard conditions (water at 20°C) (s_20,w_) using the program SEDNTERP (Schuck, 2000) to obtain standardized values. Concentration-dependent changes in the c(s) distributions for PBP2a were analyzed using isotherms based on weight-average sedimentation coefficients.

### 2.4. Production and optimization of PBP2a microcrystals

In order to optimize crystallization condition for making microcrystals for Serial Synchrotron Crystallography (SSX) and SFX experiments, we have confirmed the quality of crystals from conditions previously reported by Otero and *co-workers* (Otero *et al*., 2013). We have carried out a screening with different concentration and molecular weight of polyethylene glycols (PEGs) using the hanging-drop vapor diffusion method using 24-well plates. The best results were obtained by mixing 3 μL of protein at 15 mg/mL with 1.5 μL of precipitant solution composed of 0.1 M HEPES pH 7.0, 25% PEG 550 MME, 0.880 M NaCl and 16 mM CdCl_2_. Protein droplets were allowed to equilibrate against 1 mL of precipitant solution in the reservoir chambers at 4 °C. Large crystals appeared in about 12 h. Once the crystallization conditions for large crystals were confirmed, initial micro-crystallization trials were carried out using both the batch and the free interface diffusion (FID) methods (Kupitz *et al*., 2014). For SX experiments both at XFELs and synchrotrons, microcrystals of PBP2a were obtained on-site in the biology laboratories of these facilities by the batch with agitation method (**Figure S1**) as follows: in a 3 mL glass vial, 100 μL of the protein solution at 30 mg/mL were slowly added dropwise to 300 μL of the precipitant solution (0.1 M HEPES pH 7, 0.88 M NaCl, 25% PEG 1500, and 16 mM CdCl_2_) while agitating at 200 rpm. Upon addition of the protein, the solution turned immediately turbid and crystals of 10 μm in their longest dimensions grew at room temperature in about 1 h.

### 2.5. Initial X-ray diffraction on PBP2a microcrystals at ID30-2 at the ESRF

For this experiment, crystals of PBP2a were grown in our laboratories as shown above and then soaked in a cryoprotectant solution containing the precipitant solution supplemented with 20 % glycerol, looped, and flash-frozen in liquid N_2_. Crystals were shipped to ESRF-EBS for data collection. X-ray diffraction data collection was performed on ID30-2 beamline of the ESRF using a wavelength of 0.103 nm and a Pilatus 6M detector. Data collection was performed using a raster scan strategy. A representative diffraction pattern for PBP2a is shown in **Figure S1A**.

### 2.6. Serial synchrotron data collection on PBP2a microcrystals at ID29 at the ESRF

The SSX data were collected at ID29 of the ESRF-EBS (Orlans *et al*., 2025) using a wavelength of 1.072 Å with a pulse length of 90 μs at a repetition rate of 231.25 Hz. The X-ray beam flux at the sample was about 8 x 10^14^ ph/s and focused to 4 x 2 (H x V) μm (FWHM). Data were recorded on the JUNGFRAU 4M detector (Mozzanica *et al*., 2018) with an integration time of 95 μs to ensure full recording of the X-ray pulse and automatically corrected for pedestal and geometrical reconstruction with a LImA2 data acquisition library developed at the ESRF (https://limagroup.gitlab-pages.esrf.fr/lima2/). The sample-to-detector distance was set to 150 mm. Microcrystals of PBP2a were loaded by pipetting 3.5 μL of a crystal slurry and sandwiched between two 13 μm mylar films on a small version of the sheet-on-sheet sandwich chip (SOS chip) (Doak *et al*., 2018) as previously reported by our group (Grieco *et al*., 2024) with an opening of 5 x 10 mm.

### 2.7. Data collection, processing and structure determination of PBP2a at the EuXFEL

The SFX experiments were conducted at the SPB/SFX instrument (Mancuso *et al*., 2019) of the European XFEL (EuXFEL) during the protein crystal screening proposal ID P2967. 180 pulses were delivered in 10 Hz trains with an intra-train repetition rate of 564 kHz and 9.3 keV, with an ∼3.6 µm spot size. The average pulse energy was 2.6 mJ with a duration of 25 fs separated by 1.7 µs, and the diffraction was collected on the AGIPD 1M detector (Allahgholi *et al*., 2015, 2019). The experiment was conducted using the double-flow focusing nozzle (DFFN) liquid injector with ethanol as the sheath liquid (Oberthuer *et al*., 2017; Vakili *et al*., 2022). The flow rate was 15 to 20 µL/min for the sample and 20 µL/min for ethanol. The Helium flow rate was adjusted to reach jet speeds of about 40 m/s. Data processing was carried out off-site after the experiment. Initial hit-finding was performed using the Cheetah software (Barty *et al*., 2014) using Peakfinder 8 as algorithm (Kieffer et al., 2022). As hit finding parameters, we used an ADC threshold of 100, a minimum SNR of 5.0, and a minimum of 1 pixel per peak to identify a minimum of 7 peaks. Hits were then subjected to indexing with CrystFEL version 0.9.1 (White *et al*., 2012; White, 2019) using the algorithms MOSFLM (Powell *et al*., 2013), XDS (Kabsch, 2010), DIRAX (Duisenberg, 1992) and XGANDALF (Gevorkov *et al*., 2019). Intensities were integrated by applying integration radii of 2, 4, 6 and merged into the point group *mmm* using the CrystFEL program partialator with 1 iteration of the unity model. A representative diffraction pattern for PBP2a is shown in **Figure S1B**. Final data collection and refinement statistics are listed in **Table 1**.

**Table 1.** Data collection and refinement statistics.

| **Table 1.** Data collection and refinement statistics. |  |  |
| --- | --- | --- |
| Values for the outer shell are given in parentheses. |  |  |
|  | **LCLS-PBP2a** | **EuXFEL-PBP2a** |
| *Data collection statistics* |  |  |
| X-ray source / beamline | LCLS / MFX | EuXFEL / SPB-SFX |
| Sample delivery | MESH | DFFN |
| Sample flow rate ($\mu\text{L}/\text{min}$) | 5-10 | 15-20 |
| Photon energy (keV) | 11.2 | 9.3 |
| Repetition rate | 120 Hz | 564 kHz |
| Pulse duration (fs) | 40 | 25 |
| Beam size ($\mu\text{m}$) | 3-4 | 3.6 |
| Sample-to-detector distance (mm) | 138.7 | 121.1 |
| Temperature (K) | 293 K | 293 K |
| Detector | ePix10k | AGIPD 1MPx |
| Space group | $P2_12_12_1$ | $P2_12_12_1$ |
| a, b, c ($\text{\AA}$) | a = 81.4 $\text{\AA}$, b = 107.7 $\text{\AA}$,<br>c = 185.2 $\text{\AA}$ | a = 81.7 $\text{\AA}$, b = 106.6 $\text{\AA}$,<br>c = 186.8 $\text{\AA}$ |
| $\alpha$, $\beta$, $\gamma$ ($^\circ$) | $\alpha = \beta = \gamma = 90^\circ$ | $\alpha = \beta = \gamma = 90^\circ$ |
| Resolution range ($\text{\AA}$) | 36.0 – 2.8 (3.0-2.8) | 25.3 – 3.4 (3.5 – 3.4) |
| No. of unique reflections | 36878 (3619) | 23286 (2282) |
| Completeness (%) | 100 (100) | 100 (100) |
| Redundancy | 119 (88) | 635 (515) |
| Rsplit (%) | 25.6 (573.5) | 20.1 (409.5) |
| SNR | 2.4 (0.1) | 5.0 (0.3) |
| CC* | 0.9875 (0.4827) | 0.9950 (0.6960) |
| CC<sub>1/2</sub> | 0.9516 (0.1319) | 0.9802 (0.3196) |
| Overall B factor from Wilson plot ($\text{\AA}^2$) | 78.7 | 47.2 |
| *Refinement Statistics* |  |  |
| Resolution range ($\text{\AA}$) | 35.3 – 2.8 (2.87 – 2.8) | 25.3 – 3.5 (3.59 – 3.5) |
| No. of reflections, working set | 38413 | 20145 |
| No. of reflections, test set | 2040 | 1032 |
| R<sub>work</sub>/R<sub>free</sub> (%) | 22.5 / 29.2 | 24.2 / 31.2 |
| *No. of non-H atoms* |  |  |
| Protein | 10291 | 10291 |
| Water | 62 | 137 |
| Others | 14 | 10 |
| *R.m.s. deviations* |  |  |
| Bond length ($\text{\AA}$) | 0.005 | 0.004 |
| Bond angles ($^\circ$) | 1.400 | 1.352 |
| *Average B factors ($\text{\AA}^2$)* |  |  |
| All atoms | 105.0 | 99.0 |
| Protein | 113.9 | 111.4 |
| Water | 76.2 | 47.2 |
| *Ramachandran plot* |  |  |
| Favored (%) | 89 | 89 |
| Allowed (%) | 10 | 10 |
| Outliers (%) | 1 | 1 |
| PDB code | 9I9S | 9I9T |

All MTZ files for phasing and refinement were generated by the CTRUNCATE program from the CCP4 software package (Evans, 2011; Winn *et al*., 2011) and a fraction of 5 % reflections were included in the generated R_free_ set. Initial phases of PBP2a were obtained by molecular replacement with MOLREP (Vagin & Teplyakov, 1997) using the cryo-structure with PDB code 1VQQ (Lim & Strynadka, 2002) as the search model. The obtained model was subsequently refined using alternate cycles of automated refinement using non-crystallographic symmetry (NCS) with REFMAC5 (Murshudov *et al*., 2011, 1997) and manual inspection was performed with COOT (Emsley *et al*., 2010). The final refined structure were validated using the Protein Data Bank (PDB) validation service prior to deposition. The atomic coordinates and structure factors have been deposited in the PDB with accession code PDB 9I9T. All structure figures presented in this manuscript were generated with PYMOL version 3.0.4 (The PyMOL Molecular Graphics System, Version 3.0.4, Schrödinger, LLC, https://www.pymol.org/).

### 2.8. Data collection, processing and structure determination of PBP2a at the LCLS

The SFX data collection at the Linac Coherent Light Source (LCLS) was conducted at the Macromolecular Femtosecond Crystallography (MFX) instrument (Sierra *et al*., 2019) during protein crystal screening beamtime P10005. Diffraction snapshots were recorded on the ePix10k2M (Van Driel *et al*., 2020) detector at an X-ray energy of 11.2 keV using a pulse duration of 40 fs. The sample-to-detector distance was set to 138.7 mm. The experiment was conducted using the microfluidic electrokinetic sample holder (MESH) injector (Sierra *et al*., 2012). The MESH facilitated the continuous delivery of sample-containing microcrystals into the X-ray beam path. A customized version of OnDA (Online Data Analysis) Monitor called OM, was used for live feedback of crystal hit rate (Mariani *et al*., 2016). For the structure solution of PBP2a, a total of 1,043,234 frames were collected, of which 64,777 were classified as hits by the Cheetah software (Barty *et al*., 2014). A total of 29,212 of the identified hits (30,689 lattices) were indexed successfully with the CrystFEL program (version 0.10.1) using the following algorithms: MOSFLM (Powell *et al*., 2013) and XGANDALF (Gevorkov *et al*., 2019). The intensities were converted to structure factor amplitudes using AIMLESS from the CCP4 suite package (Winn *et al*., 2011), and a fraction of 5% reflections were included in the generated R_free_ set. Phasing was performed using molecular replacement with MOLREP (Vagin & Teplyakov, 2010) using our previously solved structure at the EuXFEL as the search model. The obtained model was refined using alternate cycles of automated refinement using non-crystallographic symmetry (NCS) with REFMAC5 (Murshudov *et al*., 2011, 1997) and manual inspection was performed with COOT (Emsley *et al*., 2010). All data collection and refinement statistics are summarized in **Table 1**. A representative diffraction pattern for PBP2a is shown in **Figure S1C**. All structure figures presented in this manuscript were generated with PYMOL version 3.0.4 (The PyMOL Molecular Graphics System, Version 3.0.4, Schrödinger, LLC, https://www.pymol.org/). The final refined structures were validated using the PDB Validation Service and submitted to the Protein Data Bank for deposition with PDB 9I9S.

## 3. RESULTS

### 3.1. Large scale production of PBP2a for SFX experiments

The original protein expression and production protocols described in (Fuda *et al*., 2004) were modified and optimized to significantly increase protein yield, addressing the demand for substantial quantities of protein required for serial crystallography experiments. To optimize induction conditions, various expression trials were performed, varying growth temperatures (37°C, 25°C, and 18°C) and IPTG concentrations (0.05 to 0.5 mM). These adjustments ensured maximal protein expression under conditions suitable for scale-up. The original five-day purification protocol, yielding 10 mg of protein per liter of culture, was condensed into a two-day protocol capable of producing up to four times as much protein. Previously, the process relied on manually packed ion exchange columns and fully manual workflows in a 4°C cold room. This included two ion exchange steps on BioRad columns (2.5 x 14 cm, 50 mL) using Macro Prep High Q and Macro Prep High S resins. The most impactful improvement was the introduction of an ammonium sulfate (AS) precipitation step. Trials with various AS saturation levels (40–80%) identified 65% as optimal (see **Figure S2A**), enabling the elimination of the first ion exchange step. Following AS precipitation, the protein was further purified using ion exchange chromatography on Macro Prep High S resin and eluted with a buffer containing 25 mM HEPES (pH 7) and 1 M NaCl (see **Figure S2B**). A major advancement was achieved with the integration of an automated purification protocol using the ÄKTA go system. This innovation significantly accelerated and simplified the process, improving both yield and reproducibility. Automation now allows for the production of up to 100 mg of highly pure protein within one week, eliminating the need for labor-intensive cold-room handling. To ensure the highest purity, a size-exclusion chromatography (SEC) step was added to the workflow (see **Figure S2C**). This additional step enhanced protein quality without sacrificing efficiency, resulting in a streamlined and robust protocol suitable for high-throughput applications. A detailed schematic of the purification workflow is shown in **Figure S3**.

### 3.2. Homogeneous and bulk production of PBP2a microcrystals for SFX experiments

To optimize crystallization conditions for producing microcrystals suitable for SSX and SFX experiments, we first tested the crystallization parameters previously reported by Otero *et al*. (Otero *et al*., 2013) to verify the quality of the resulting crystals. A screening was then conducted to evaluate the effects of varying polyethylene glycol (PEG) concentrations and molecular weights, using the hanging-drop vapor diffusion method in 24-well plates. The optimal conditions were achieved by mixing 3 μL of protein at 15 mg/mL with 1.5 μL of a precipitant solution containing 0.1 M HEPES (pH 7.0), 25% PEG 550 MME, 0.880 M NaCl, and 16 mM CdCl_2_. The drops were equilibrated against 1 mL of precipitant solution in the reservoir at 4°C.

After confirming conditions for growing large crystals and assessing their diffraction quality, we initiated micro-crystallization trials using the batch and the free interface diffusion (FID) methods (Kupitz *et al*., 2014). These trials explored a range of variables, including: 1) PEG concentration: 20-45%; 2) PEG type: PEG 550 MME, PEG 1000, PEG 1500; 3) salt concentration: 100-880 mM NaCl; 3) protein-to-precipitant ratios: 1:1, 1:2, 1:3; protein concentrations: 20-35 mg/mL; and 4) temperature: 18°C and 4°C. The FID method provided the best results under the following conditions: 0.1 M HEPES (pH 7.0), 25% PEG 1500, 0.880 M NaCl, 16 mM CdCl_2_, a protein concentration of 30 mg/mL, and a 1:1 protein-to-precipitant ratio at 18°C. Upon identifying optimal conditions, efforts were directed toward scaling up microcrystal production. Using a batch crystallization system with continuous agitation, microcrystals were reproducibly obtained in large quantities (**Figure S4A**). This was achieved by mixing 100 μL of protein at 30 mg/mL with 300 μL of precipitant solution at a 1:3 protein-to-precipitant ratio. Best microcrystals are shown in **Figure S4B**.

### 3.3. X-ray diffraction of PBP2a microcrystals at the synchrotron

Before even performing a TR-SFX experiment, a couple of steps must be taken to confirm that the sample is suitable for this type of experiment. First, it is necessary to confirm that the crystals are of good diffracting quality. Second, it is desirable to solve the protein structure in its apo form using, for example, serial synchrotron crystallography (SSX). To this end, once microcrystals of PBP2a were optimized, we measured them in a classic X-ray crystallography experiment to check for their diffraction quality. Microcrystals grown as described in section 3.2 in our labs, were looped, flash frozen and shipped to ID30-2 at ESRF-EBS for X-ray diffraction. Data collection was performed using a raster scan strategy. A total of 5,000 frames were collected from one loop, from which 500 were successfully indexed with CrystFEL. Although the number of images collected was not sufficient to allow us to determine the structure of PBP2a, which in fact was not the purpose of this quality test, the experiment allowed us to confirm the high quality of our microcrystals of which diffraction up to 2.2 Å was observed (**Figure S1A**). Then we moved forward to the next step. For the serial synchrotron crystallography (SSX) experiments at the ID29 beamline of ESRF-EBS, frozen protein was shipped to ESRF-EBS and microcrystals of PBP2a were prepared at ID29 labs as shown in section 3.2 and subsequently loaded into a small SOS chip as previously described (Grieco *et al*., 2024). The mylar-sample-mylar sandwich was placed between the two half frames of the chip holder leading to further spreading and thinning of the film by capillary action. The two frame halves were then sealed together with one clamping screw. This chip design can ensure optimal crystal visualization and minimized background noise during data collection. Serial data collection at ID29 was conducted as previously reported by our group (Grieco *et al*., 2024). The experimental setup is shown in **Figure 1**. Unfortunately, just a few diffraction images were observed containing few diffraction spots. Indexing with CrystFEL was attempted unsuccessfully with a low indexing rate that did not allow us to solve the structure of PBP2a.

**Figure 1.**
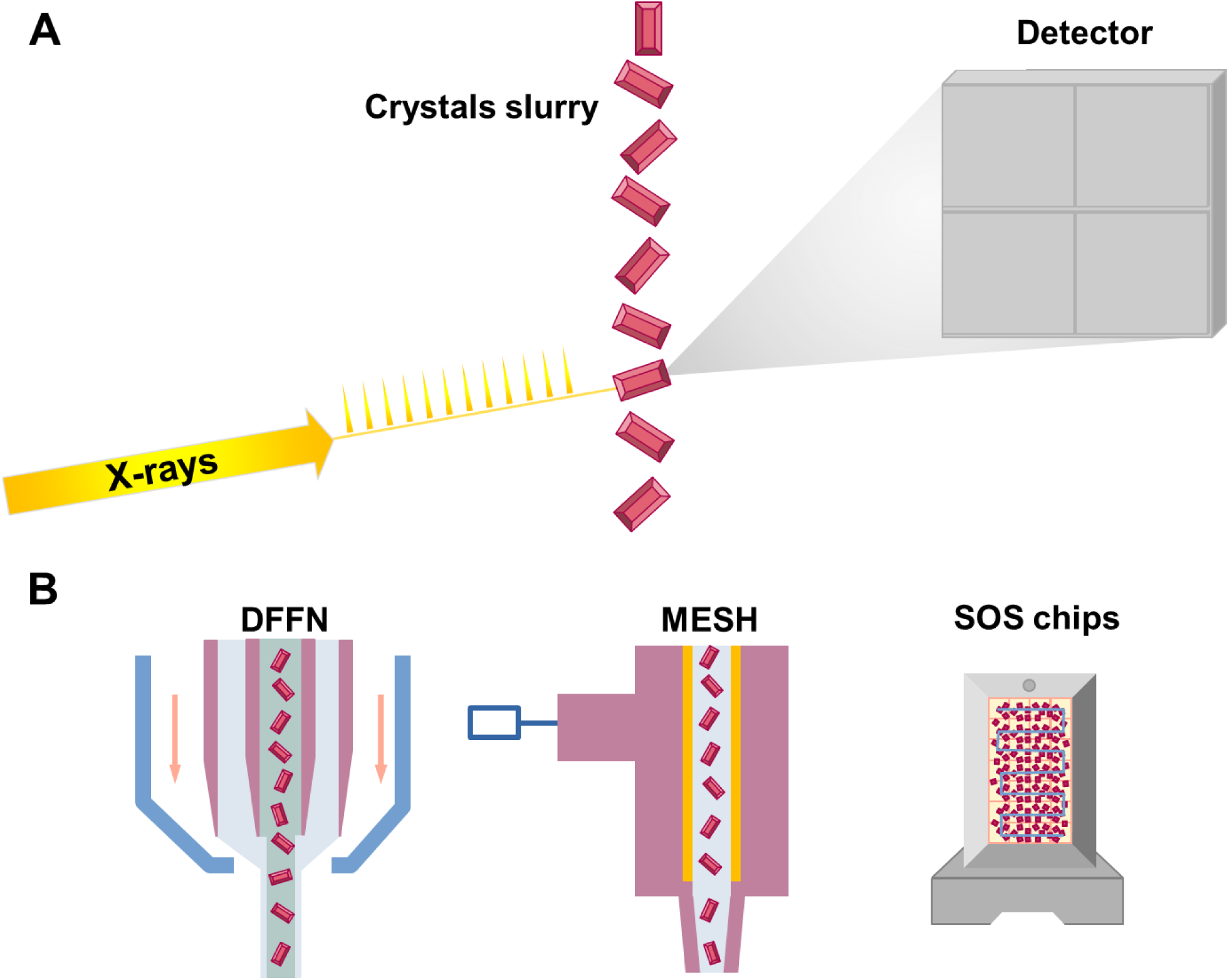
Experimental setup and sample delivery devices used for the serial crystallography experiments presented in this study. **A)** Schematic of the serial crystallography setup illustrating the X-ray source delivering X-ray pulses from XFELs and synchrotrons to a stream of randomly oriented microcrystals delivered in a serial fashion. Each crystal produces a single diffraction pattern. The scattered X-rays are recorded by a high-speed detector. **B)** Delivery devices employed during our experiments. The double-flow focusing nozzle (DFFN) (Oberthuer et al., 2017), used at the SPB/SFX instrument, delivers a stable, continuous stream of microcrystals into the X-ray beam, ensuring efficient interaction and minimal sample waste. The MESH device (Sierra et al., 2012), employed at the MFX instrument, uses electrokinetic forces to precisely control the flow of microcrystals in a liquid medium. The sheet-on-sheet (SOS) chip (Doak et al., 2018), used at beamline ID29, sandwiches microcrystals between thin mylar sheets, providing a stable and consistent delivery system for synchrotron-based serial data collection.

### 3.4. SFX data collection on PBP2a microcrystals at XFELs

SFX experiments at the European XFEL were conducted at the SPB/SFX instrument during the protein crystal screening experiment P2967. We used the DFFN injector (Oberthuer *et al*., 2017), specifically designed for continuous sample delivery into the XFEL beam. The DFFN is a Gas Dynamic Virtual Nozzle (GDVN) with an additional liquid line resulting in an inner jet of sample and an outer jet of ethanol. Data from microcrystals of PBP2a were successfully collected and a representative diffraction pattern is shown in **Figure S1B**. During this experiment, the hit rate varied between 1.18 and 3.66 %. All data collection statistics are listed in **Table 1**. At LCLS, SFX data collection was conducted at the MFX instrument (Sierra *et al*., 2019) during the protein crystal screening experiment P10005. In this case, we used the MESH injector (Sierra *et al*., 2012). This injector, based on microfluidics technology, facilitated the continuous and stable delivery of PBP2a microcrystals into the XFEL beam path. The MESH injector employed syringe pumps or pressure-based systems to precisely control the flow of crystal-containing solution, optimizing crystal injection rates and minimizing sample wastage. During our experiment the hit rate varied from 1 to 24 %. Data from microcrystals of PBP2a were successfully collected and a representative diffraction pattern is shown in **Figure S1C**. The experimental setups used in these two XFEL experiments are shown in **Figure 1**. All data collection statistics are listed in **Table 1**.

### 3.5. Room temperature crystal structures of PBP2a

We have successfully determined the structure of PBP2a using data collected exclusively from SFX experiments at both the EuXFEL and the LCLS facilities. All data collection statistics are reported in **Table 1**. PBP2a microcrystals belonged to the space group P2_1_2_1_2_1_ with the following unit cell dimension a= 81.7 Å, b=106.6 Å, c=186.8 Å, α= β= γ= 90° and diffracted to a maximum resolution of 3.4 Å resolution. To solve the structure of PBP2a from data collected at the EuXFEL, a total of 3,136,404 frames were collected, of which 87,337 were classified as hits by CrystFEL. About 35% of the identified hits could then be indexed, giving rise to a total of 30,689 indexed, integrated, and merged lattices. The structure was solved by molecular replacement using the PDB entry 1VQQ (Lim & Strynadka, 2002) as a search model without water, and ion molecules. The structure was refined at a final resolution of 3.5 Å with R_work_ and R_free_ of 24.2 % and 31.2%, respectively. In the case of the data collected at the LCLS, the microcrystals of PBP2a also belonged to the space group P2_1_2_1_2_1_ with the following unit cell dimensions a= 81.4 Å, b=107.7 Å, c=185.2 Å, α= β= γ= 90° and diffracted to 2.5 Å resolution. To solve the structure of PBP2a from data collected at LCLS, a total of 1,043,234 frames were collected, of which 64,777 were classified as hits by Cheetah (Barty *et al*., 2014). About 45.1 % of the identified hits could then be indexed, leading to a total of 29,212 indexed, integrated, and merged lattices. The structure of PBP2a was solved by molecular replacement using our previous SFX structure solved at the EuXFEL as a search model without water, and ion molecules. The structure was refined at a final resolution of 2.8 Å with R_work_ and R_free_ of 22.5 % and 29.2 %, respectively. All data collection and refinement statistics are listed in **Table 1**.

Unlike previously observed in the cryogenic structure of PBP2a (Lim & Strynadka, 2002), the two room temperature structures presented here could be modeled fully in the electron density from N-terminus to the C-terminus, filling all the residues without any interruptions in the case of PBP2a from LCLS data. The resulting experimental maps were of good quality revealing the presence of solvent ions, such as five chloride ions and seven cadmium ions in the PBP2a structure from the EuXFEL, and seven chloride ions and seven cadmium ions in the PBP2a structure from the LCLS data. Our structures also revealed the presence of a high number of water molecules: 135 water molecules in our PBP2a structure from EuXFEL and 82 water molecules in the PBP2a structure from LCLS. The quality of our PBP2a structures can be assessed from the electron 2mFo-DFc density maps shown for newly modelled regions (88-93, 504-507 and 605-610), the allosteric and catalytic sites, as well as for the cadmium ions sites (**Figures 2 and S7**). All our structures revealed the presence of two PBP2a protein molecules in the asymmetric unit (**Figure S5**). Due to the conformational changes and the allosteric behavior previously described for this enzyme, we will consider both molecules, A & B, in our discussion.

**Figure 2.**
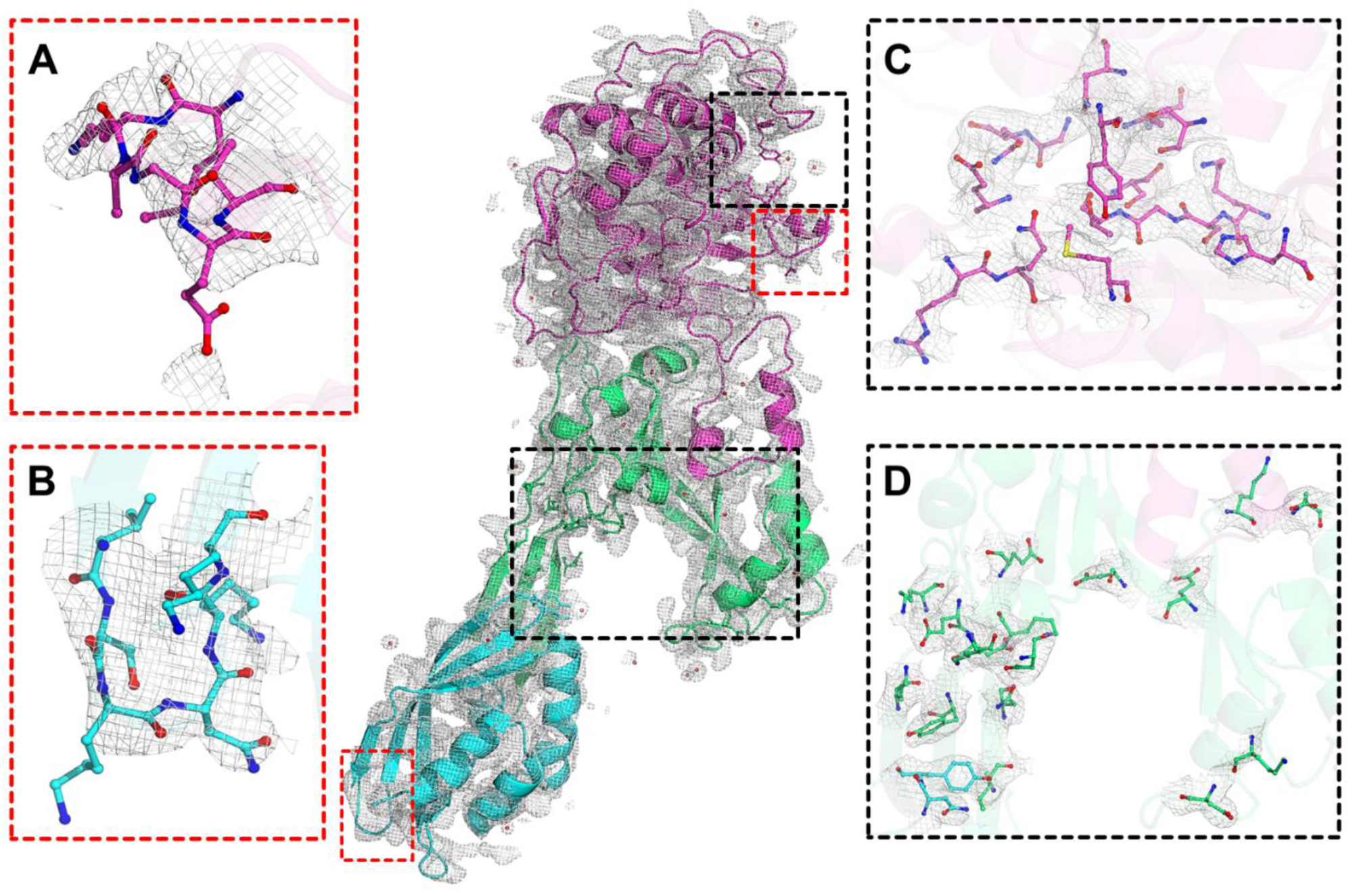
Room temperature structure of PBP2a at XFELs. The central image displays the 2mF_o_-DF_c_ electron density maps contoured at 1σ around chain A of the room-temperature PBP2a structure solved at LCLS. The structure is depicted in cartoon representation, with the N-terminal extension domain (residues 27–138) colored cyan, the allosteric domain comprising lobes 1–3 (residues 139–326) in green, and the transpeptidase domain (residues 327–668) in violet. Panels A–D on either side of the structure highlight specific features: newly modeled regions 605–610 (panel A) and 88–93 (panel B), as well as the catalytic site (panel C) and the allosteric site (panel D).

### 3.6. Structural Comparison of apo PBP2a with related cryogenic structures

A comparison of our room temperature structures to other PBP2a structures determined in cryogenic conditions reveals notable differences in the unit cell dimensions. As shown in **Table 1**, the two structures of the apo PBP2a presented in this study exhibit slight changes in the unit cell parameters; however, display an important reduction or shrinking of the unit cell dimensions of up to 7 Å in *b* dimension compared to those from the cryo-structure in its apo form (PDB 1VQQ, (Lim & Strynadka, 2002)), which showed a unit cell dimensions of a= 80.9 Å, b=100.6 Å, c=186.2 Å, α= β= γ= 90°. These differences in unit cell dimensions are evident when comparing the packing of the PBP2a room temperature structures with that observed under cryogenic conditions (**Figure S6**). This variation in unit cell dimensions is a consequence of the increased flexibility and natural conformational states that, overall, proteins adopt at near physiological temperatures. Such differences in unit cell dimensions are indicative that cryogenic conditions may impose constraints on the protein structure and potentially mask biologically relevant conformational states.

Based on our experience with highly dynamic and allosteric proteins (Doppler *et al*., 2023; Grieco *et al*., 2023, 2024), we emphasize the importance of including all protein molecules in the asymmetric unit (ASU) in the analysis. In this manuscript, we analyze both PBP2a molecules in the ASU, rather than just one, to account for their conformational changes and allosteric behavior. In this regard, the two molecules of the SFX structures of the apo PBP2a presented in this study were further analyzed by comparing them with the structure of the apo PBP2a previously reported at cryogenic temperature (PDB 1VQQ, (Lim & Strynadka, 2002)). A superimposition of all molecules in our SFX structures is shown in **Figure S7A** and the superimposition of the SFX structures to those from the cryo-structure is shown in **Figure S7B**. Overall, all apo PBP2a structures aligned very well with each other with subtle differences found across the protein showing RMSD values for the Cα atoms between 0.369 and 0.811 Å and with an average value of 0.590 Å. The RMSD values for all atoms were between 0.643 and 1.320 Å and with an average value of 0.981 Å. However, as the crystal structures are obtained at room temperature, notable differences in certain structural elements of the transpeptidase and allosteric domains have been observed (**Figure 3**): 1) β-strands 1-5 at the transpeptidase domain are generally shorter than those in the cryo-structure, with β-3 showing increased flexibility; 2) the final turn of the C-terminal region of α-9 is shorter compared to the cryo-structure; 3) the helix α1-np at lobe 1; 4) the helix α5-np at lobe 2; and 5) helix α-4 and β-strands β-2a and β-2b transition into loops at the transpeptidase domain. Despite the above differences, the cryo-structure contains three missing regions (88-93, 504-507 and 605-610) that have been completely modelled in our structures (**Figure 2**). Of special interest is the region 605-610, in which the three residues Glu609, Thr610 and Gly611 form a short 3_10_-helix. This region 605-610 is not even seen in the chain A of complex of PBP2a with ceftaroline; however, is fully modelled in the chain B.

**Figure 3.**
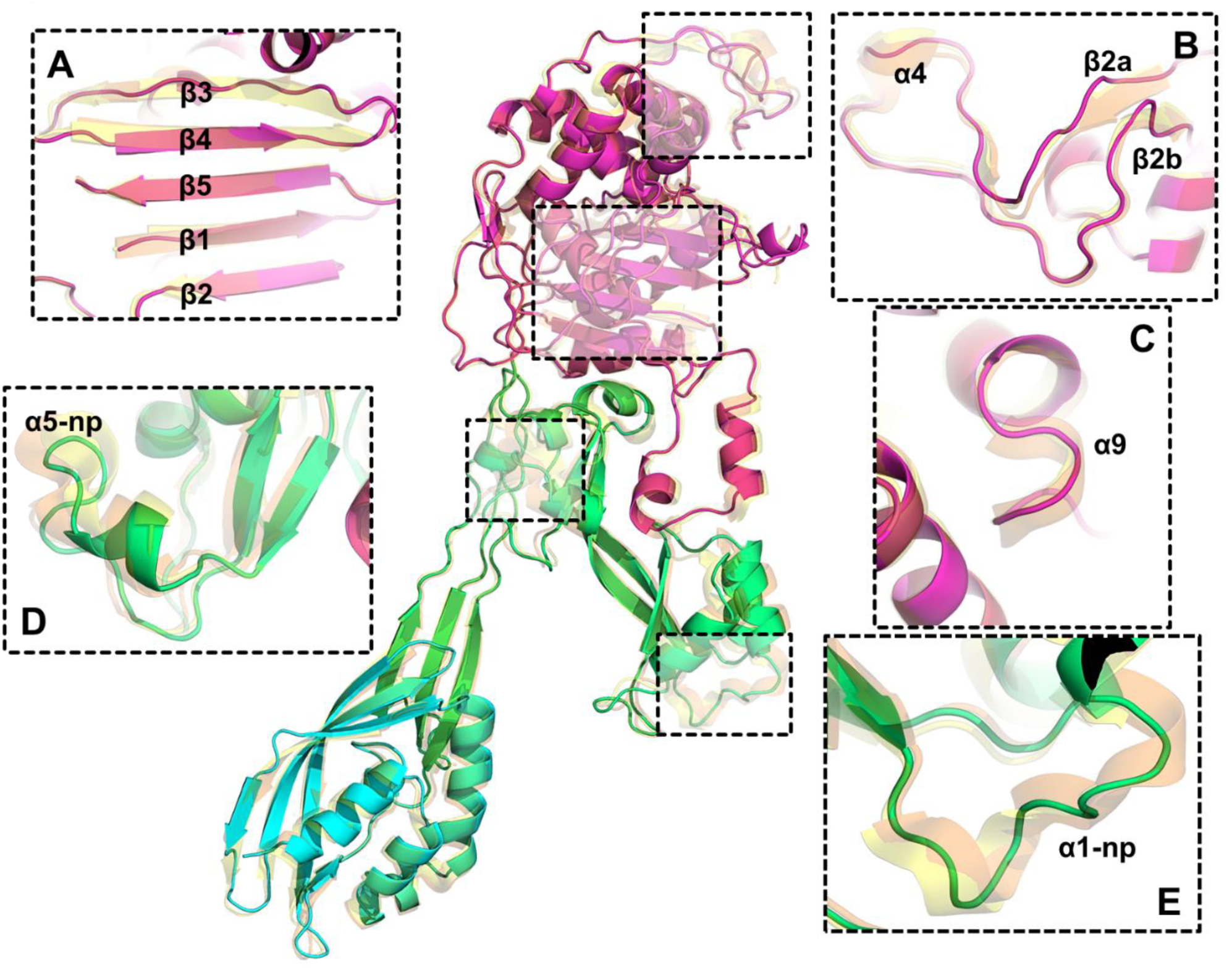
Structural differences between room-temperature and cryogenic PBP2a crystal structures. The central image displays a superimposition of chain A of the room-temperature PBP2a structure solved at LCLS to chains A and B of the apo PBP2a (PDB 1VQQ) (Lim & Strynadka, 2002). The structure is depicted in cartoon representation with the same color code as in Figure 2. Panels A–E on either side of the structure highlight this most notable differences: **A)** the β-Strands 1–5 in the transpeptidase domain; **B)** the helix α-4 and the β-strands β-2a and β-2b in the transpeptidase domain; **C)** the final turn of the C-terminal region of α-9 is truncated; **D)** the helix α5-np at lobe 2; and **E)** the helix α1-np at lobe 1.

To develop a further comparison of our room temperature structures with the cryogenic ones, we conducted a detailed interaction pattern analysis of PBP2a structures using ARPEGGIO (Jubb *et al*., 2017) (**Table S1**). Comparing the room-temperature XFEL structures of apo PBP2a reveals subtle differences. The total contacts are slightly higher in the LCLS structure (7,331 for Chain A and 7,191 for Chain B) than in the EuXFEL structure (6,688 for Chain A and 6,540 for Chain B), suggesting a marginally more extensive interaction network in LCLS. Van der Waals and polar contacts follow a similar trend, with the structure obtained at LCLS having slightly more interactions (**Table S1**). However, water-mediated polar contacts and hydrogen bonds are slightly higher in the EuXFEL structure, indicating differences in hydration or water molecule placement (**Table S1**). Salt bridge counts are higher in the LCLS data for both chains, with Chain A having 78 (compared to 71 in EuXFEL) and Chain B having 83 (compared to 43 in EuXFEL).

In contrast, the cryogenic structure of apo PBP2a (PDB 1VQQ, (Lim & Strynadka, 2002)) and the ligand-bound structure with ceftaroline (PDB 3ZFZ, (Otero *et al*., 2013)) exhibit significantly denser interaction networks compared to the XFEL structures (**Table S1**). These cryogenic structures have higher totals for contacts, Van der Waals interactions, polar contacts, and hydrogen bonds, which can be attributed to the combined effects of higher resolution capturing finer details and the stabilizing influence of low temperatures reducing conformational variability. In contrast, the room temperature structures reflect the protein’s dynamic and flexible nature under near-physiological conditions, resulting in less densely populated interaction networks. For example, the chains A of PBP2a in 1VQQ and 3ZFZ contain 10,500 and 10,017 contacts, respectively, both substantially higher than the totals for the room temperature structures. Water-mediated interactions are particularly abundant in 1VQQ and 3ZFZ, with polar contacts exceeding 580 in both, underscoring the role of cryogenic conditions and ligand binding in stabilizing interactions (**Table S1**). Meanwhile, the XFEL structures likely capture more dynamic substates, emphasizing the inherent flexibility and conformational variability of apo PBP2a.

### 3.7. Structural changes in the allosteric site of PBP2a

The allosteric site, a large cavity with an accessible surface area of 7,983.5 Å^2^, remains structurally unchanged in terms of residue conformation upon antibiotic or peptidoglycan binding (Otero *et al*., 2013). However, ligand binding induces localized changes that propagate throughout the protein, ultimately reaching the catalytic site (Otero *et al*., 2013). This structurally intricate allosteric region comprises residues 166-240 (lobe 1), 258-277 (lobe 2), 364-390 (lobe 3), and the upper portion of the N-terminal domain. It features a complex network of hydrogen bonds and salt bridges, essential for maintaining structural integrity and facilitating ligand-driven transitions (**Figures 4 and S8, and Table S2**). A comparative analysis across room-temperature XFEL and cryogenic structures reveals conserved interactions, such as Lys148-Asp295, Lys316-Glu294, Lys319-Glu294, and Lys316-Tyr297, which persist regardless of antibiotic binding (**Table S2**). These interactions do not directly influence ligand binding but seem to be vital for stabilizing the allosteric site. However, the binding of ceftaroline triggers notable structural rearrangements, including the formation of new salt bridges (Lys198-Asp202 and Lys219-Asp221) and disruption of others, such as Lys273-Glu94. Interestingly, this disrupted interaction is also absent in the cryogenic apo structure (PDB 1VQQ) (Lim & Strynadka, 2002) and molecule B of PDB 3ZFZ (Otero *et al*., 2013). Moreover, the adaptive response of the protein to ligand interaction is further highlighted by conformational changes in lobe 1, where new salt bridges replace pre-existing hydrogen bonds.

**Figure 4.**
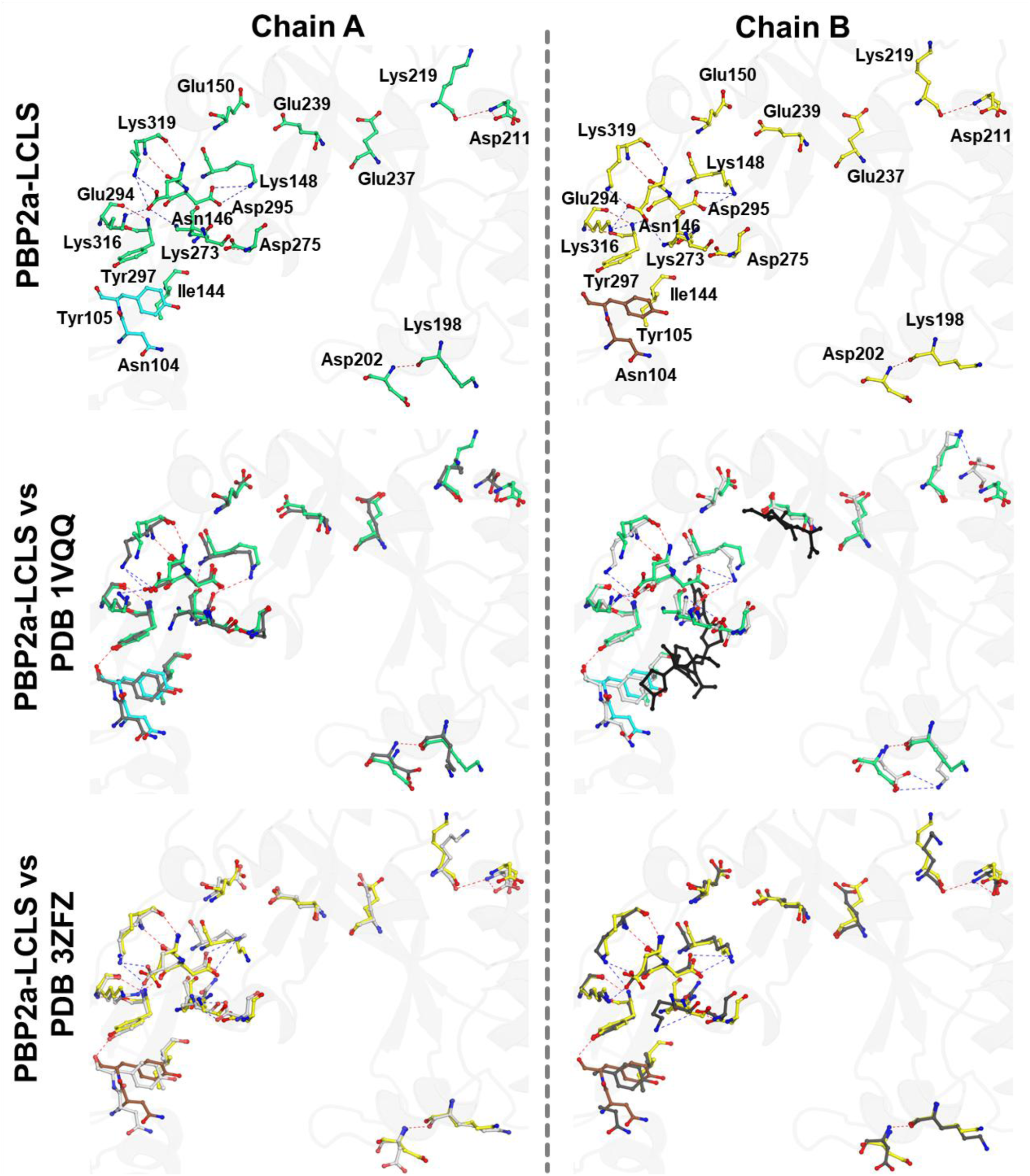
Structural alterations at the allosteric site of PBP2a in room-temperature structure PBP2a-LCLS compared to related structures. Upper panel depict all residues forming the allosteric site of PBP2a for chains A and B of the room-temperature PBP2a-LCLS structures. Hydrogen bonds within the allosteric site are highlighted in red, while salt bridges are shown in blue. Middle panel depict the superposition of the allosteric site of PBP2a-LCLS with that of the apo structure (PDB 1VQQ) (Lim & Strynadka, 2002). To illustrate the differences in the interaction network with respect to the PBP2a-LCLS, only the interactions in (PDB: 1VQQ) (Lim & Strynadka, 2002) are shown using the same color code (hydrogen bonds in red and salt bridges in blue). Lower panel depicts the superposition of the allosteric site of PBP2a-LCLS with that of the PBP2a in complex with ceftaroline (PDB 3ZFZ) (Otero et al., 2013). To illustrate the differences in the interaction network with respect to the PBP2a-LCLS, only the interactions in (PDB 3ZFZ) (Otero et al., 2013) are shown using the same color code (hydrogen bonds in red and salt bridges in blue). Both ceftaroline and muramic acid ligands are shown as black sticks. For clarity, only residues in the upper panel have been labeled.

A detailed examination of the lysine residues within this network reveals their critical roles in mediating structural stability and ligand-induced transitions. Lys148, Lys198, Lys219, Lys273, and Lys319 exhibit higher flexibility, adopting distinct conformations across the analyzed structures (**Figures 4 and S8**). Notably, Lys148 forms a stable salt bridge with Asp295 in room-temperature XFEL structures, with bond lengths ranging from 3.8 to 4.3 Å, absent in the cryogenic apo structure (**Table S2**). Upon ceftaroline binding, this interaction strengthens (3.1-3.9 Å), indicating its role in stabilizing the ligand-binding conformation. Additionally, Lys148 forms hydrogen bonds with Asn146 in the ligand-bound state, emphasizing its dynamic contribution to ligand recognition (**Figures 4 and S8, and Table S2**). Similarly, Lys198 consistently interacts with Asp202 through hydrogen bonds across all structures, with bond lengths of 3.0-3.4 Å. In room-temperature structures, these interactions display minor variations, reflecting increased flexibility (**Table S2**). Upon ceftaroline binding, Lys198 forms additional salt bridges with Asp202 (3.3-4.3 Å), absent in apo forms, suggesting its involvement in stabilizing the ligand-induced conformation and mediating communication between the allosteric and catalytic sites (**Figures 4 and S8, and Table S2**). Lys219 also demonstrates a dual role, interacting with Asp221 via hydrogen bonds in room-temperature structures (3.8-3.9 Å) (**Table S2**). These interactions are absent in the cryogenic apo structure, suggesting that room-temperature conditions stabilize conformations that enable such interactions. Interestingly, although far from the binding site, the binding of ceftaroline brakes the hydrogen bond interaction established between residues Tyr223 and Glu189, resulting in a significant conformational change in which the helix α3np at lobe 1, which is shifted approximately 6 Å relative to the cryo-structure, so that there is a new strong salt bridge forming between Lys219 and Asp221, highlighting its dual role in maintaining structural integrity in both the apo and ligand-bound states (**Figure 5A**). The network involving Lys219 and Asp221 may act as a hinge point for conformational transitions triggered by ligand binding. Lys273 participates in distinct interaction networks, forming a salt bridge with Glu294 in room-temperature structures (3.4-3.6 Å), which shortens compared to cryogenic structures (4.2 Å), indicating increased stabilization (**Table S2**). However, this interaction is disrupted in the ligand-bound state as Glu294 interacts directly with ceftaroline. Lys273 compensates by forming a new salt bridge with Asp275 (4.4-4.5 Å), stabilizing the antibiotic-bound conformation and facilitating structural reorganization (**Table S2**). Finally, Lys319 bridges the allosteric site with its surroundings, forming strong salt bridges with Glu294 (2.9-3.6 Å) and hydrogen bonds with Tyr297 (2.7-3.4 Å). While partially present in cryogenic structures, these interactions are more stable in room-temperature structures (**Table S2**). In the ceftaroline-bound state, Lys319 disrupts its interaction with Glu294, adapting to ligand binding and preserving allosteric site integrity during dynamic transitions (**Table S2**).

**Figure 5.**
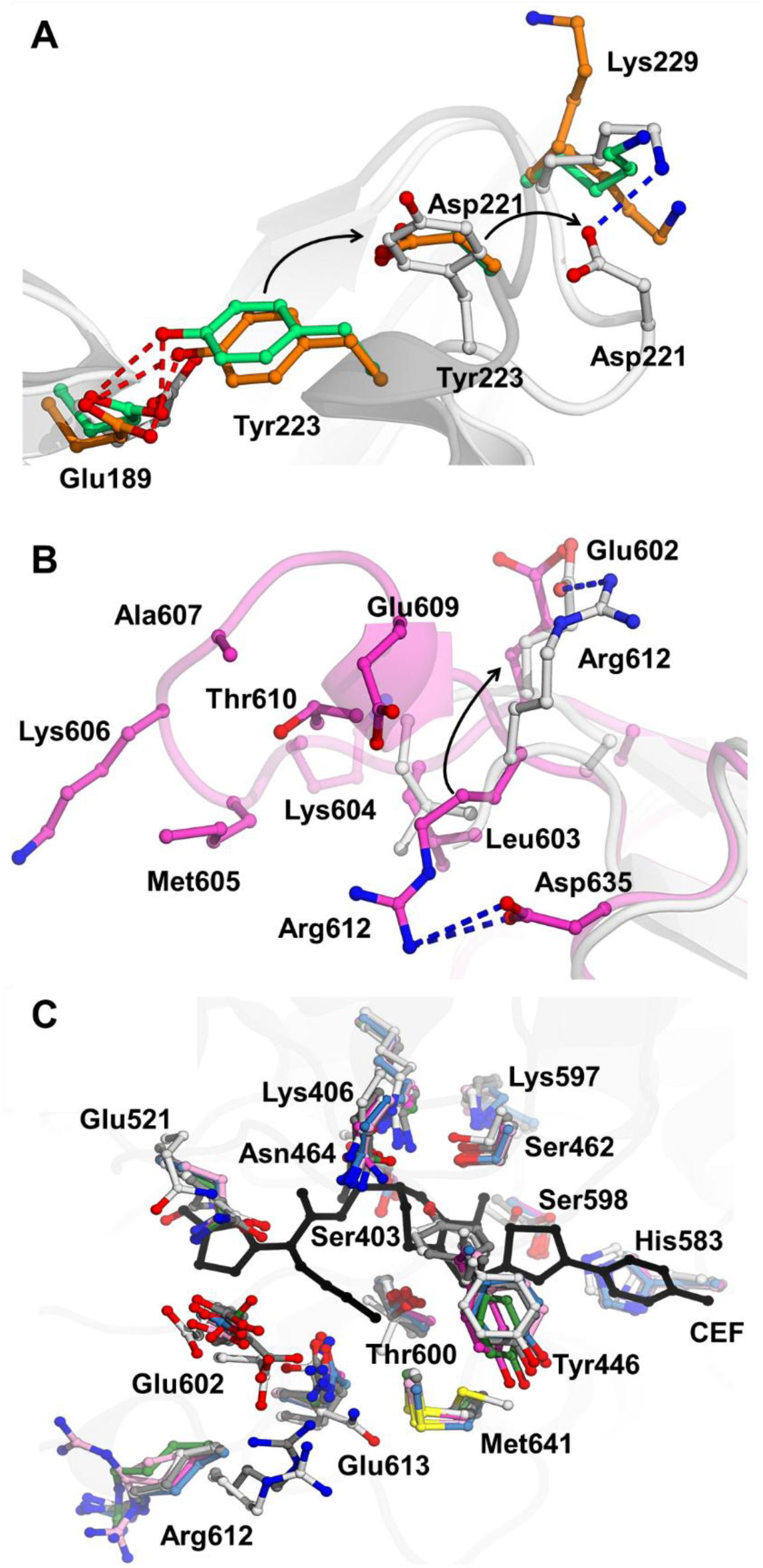
Ligand-induced conformational changes. **A)** Close-up view of helix α3np in lobe 1, comparing the room-temperature structures (PBP2a-LCLS in green and PBP2a-EuXFEL in orange) with the PBP2a-ceftaroline complex structure (PDB 3ZFZ in white, (Otero et al., 2013)). The figure highlights the disruption of the Tyr1223-Asp221 hydrogen bond (red dashed line) observed in the room-temperature structures and the formation of the Lys1229-Asp221 salt bridge (blue dashed line) upon ceftaroline binding. Key residues are depicted as sticks. **B)** Close-up view of the significant conformational change in Arg612 upon ceftaroline binding. The figure shows the disruption of the Arg612-Asp635 interaction and the formation of a new Arg612-Asp605 interaction. Key residues are depicted as sticks. **C)** Close-up of the catalytic site in chains A and B, comparing the room-temperature structures presented in this study (PBP2a-LCLS in magenta and PBP2a-EuXFEL in orange), the apo PBP2a cryo-structure (PDB 1VQQ in grey, (Lim & Strynadka, 2002)), and the PBP2a-ceftaroline complex structure (PDB 3ZFZ in white, (Otero et al., 2013)). The inward- and outward-facing conformations of Tyr446 at the catalytic site are highlighted. All residues in the catalytic site are depicted as sticks.

Together, this intricate network of lysine-mediated interactions underscores the adaptive flexibility and stability of the allosteric site in PBP2a. Lys148 and Lys319 are pivotal in modulating interactions with Asp295 and Glu294, respectively, while Lys198, Lys219, and Lys273 stabilize the apo state and facilitate ligand-induced structural transitions. The shorter interaction distances observed at room temperature suggest enhanced dynamics and adaptability, essential for ligand recognition and binding.

### 3.8. Structural changes in the catalytic site of PBP2a

Analysis of room-temperature XFEL structures of PBP2a provided significant insights into the enzyme’s catalytic site interactions and conformational dynamics, particularly in comparison to previously reported cryo-structures of apo PBP2a (PDB 1VQQ, (Lim & Strynadka, 2002)) and the ceftaroline-bound complex (PDB 3ZFZ, (Otero *et al*., 2013)). While the catalytic site exhibited a broadly conserved interaction pattern across all datasets, unique features and differences were evident under room-temperature conditions, underscoring the impact of temperature on hydrogen bonding and flexibility within the catalytic pocket (**Table S3**). In this regard, in the room-temperature structures, the nucleophile Ser403 formed consistent hydrogen bonds with Lys406, Lys597, and Thr600. Notably, the Ser403-Thr600 interaction was observed at 2.8 Å in both XFEL structures but was mostly absent in cryogenic structures (**Figures 6 and S9, and Table S3**). Similarly, Lys406 maintained interactions with Ser462 (3.8 Å in XFEL structures) and Asn464 (2.8-3.3 Å across all datasets) (**Table S3**). These findings suggest that room-temperature conditions preserve a dynamic interaction network closer to physiological states, while cryogenic cooling restricts the conformational sampling of side chains.

**Figure 6.**
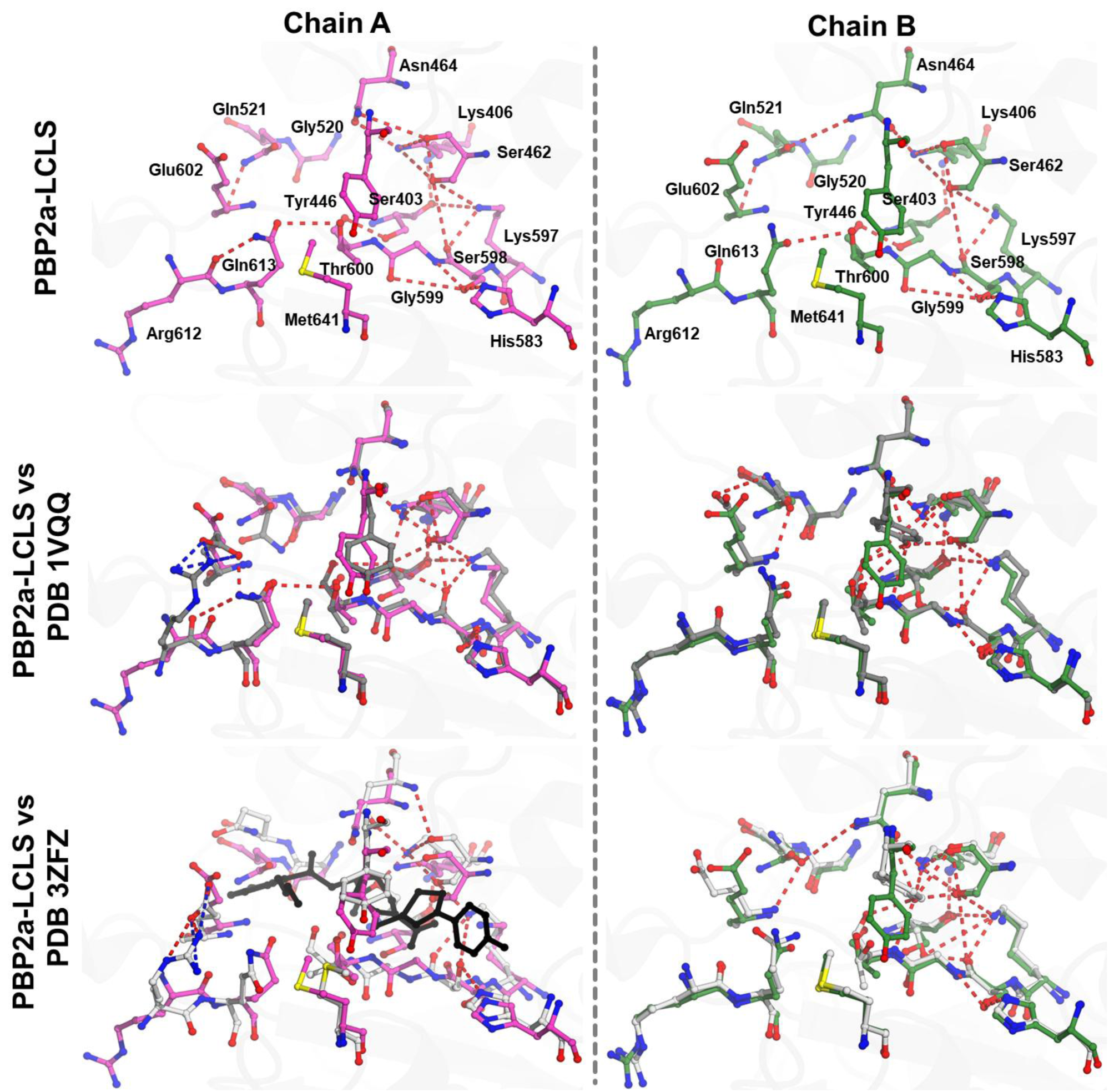
Structural alterations at the catalytic site of PBP2a in room-temperature structure PBP2a-LCLS compared to related structures. Upper panel depict all residues forming the catalytic site of PBP2a for chains A and B of the room-temperature PBP2a-LCLS structures. Hydrogen bonds within the catalytic site are highlighted in red, while salt bridges are shown in blue. Middle panel depicts the superposition of the catalytic site of PBP2a-LCLS with that of the apo structure (PDB 1VQQ) (Lim & Strynadka, 2002). To illustrate the differences in the interaction network with respect to the PBP2a-LCLS, only the interactions in (PDB: 1VQQ) (Lim & Strynadka, 2002) are shown using the same color code (hydrogen bonds in red and salt bridges in blue). Lower panel depict the superposition of the catalytic site of PBP2a-LCLS with that of the PBP2a in complex with ceftaroline (PDB 3ZFZ) (Otero et al., 2013). To illustrate the differences in the interaction network with respect to the PBP2a-LCLS, only the interactions in (PDB 3ZFZ) (Otero et al., 2013) are shown using the same color code (hydrogen bonds in red and salt bridges in blue). Ceftaroline ligand is shown as black sticks. For clarity, only residues in the upper panel have been labeled.

Binding of ceftaroline in chain A of PDB 3ZFZ induced significant conformational changes within the catalytic site. For instance, ceftaroline binding disrupted Ser403’s interactions with Lys597 and Thr600, while retaining its bond with Lys406 and establishing new interactions with the antibiotic (**Figures 6 and S9, and Table S3**). Ligand-induced perturbations also disrupted several key hydrogen bonds and salt bridges, including those between Asn521-Glu602, Ser598-Ser599, Ser598-Gly599, and Thr600-Gln613 (**Table S3**). However, conserved interactions, such as His583-Ser598, Lys406-Asn464, Lys406-Ser462, and Lys597-Ser598, were maintained across all structures, emphasizing their critical role in stabilizing the catalytic site under varying conditions (**Figures 6 and S9, and Table S3**). Unique interactions in the ceftaroline complex include a hydrogen bond between Lys406 and Tyr446 (2.9 Å), which was absent in apo forms, and salt bridges involving Glu602 and Arg612 (2.8-4.0 Å) (**Table S3**). These specific interactions likely stabilize the antibiotic in the catalytic site, contributing to its inhibitory mechanism. Structural differences in the catalytic site were further highlighted by the identification of a previously unresolved loop near the active site forming a 3_10_-helix in the room-temperature structures (**Figure 2**). This structural element, though not fully visible in the PBP2a-ceftaroline complex, undergoes a conformational change when the catalytic site opens to accommodate the antibiotic, causing the salt bridges between Arg612 and Asp635 to break and a new salt bridge to form with Glu602 (**Table S3** and **Figure 5B**). This observation, absent in prior structures of apo PBP2a, highlights the value of room-temperature crystallography for capturing dynamic features essential for understanding enzyme function. Additionally, residue Tyr446 exhibited temperature-dependent conformations, adopting an outward-facing conformation in the room-temperature structures, while cryogenic conditions favored an inward-facing conformation in certain chains, suggesting temperature-dependent stabilization (**Figure 5C**).

Together, these results highlight the dynamic nature of PBP2a’s catalytic site and its responsiveness to antibiotic binding. The unique interaction patterns and structural elements captured under room-temperature conditions provide deeper insights into the enzyme’s functional mechanisms, interaction landscape, and the influence of temperature on structural dynamics, which were previously obscured in cryogenic studies.

### 3.9. Structural changes connecting allosteric and catalytic sites

As initially described by Otero et al. (Otero *et al*., 2013) and explored in subsequent studies (Fishovitz *et al*., 2014; Mahasenan *et al*., 2017), a complex network of salt bridges extending from the allosteric site to the catalytic site of PBP2a plays a pivotal role in its regulation. The binding of the antibiotic ceftaroline induces a reorganization of these salt bridges, connecting the two sites in a manner that results in the opening of the catalytic site (Fishovitz *et al*., 2014; Mahasenan *et al*., 2017; Otero *et al*., 2013). This structural rearrangement is essential for exposing Ser403, a residue critical for the transpeptidase reaction. In the present study, we analyzed the salt bridge interactions in two newly determined room-temperature XFEL structures of PBP2a and compared them with cryogenic structures of apo PBP2a (PDB 1VQQ) (Lim & Strynadka, 2002) and its complexes with ceftaroline (PDB 3ZFZ) and peptidoglycan (PDB 3ZG5) (Otero *et al*., 2013). By examining these structures under different conditions, we aim to understand the potential role of salt bridges in modulating structural dynamics and allosteric communication. This comprehensive comparison reveals both conserved and variable salt bridge interactions, offering insights into the influence of temperature-dependent dynamics and ligand-induced conformational states.

Our analyses identified a total of 18 salt bridges that are conserved across all structures (**Tables S5 and S6**). These conserved interactions, illustrated in **Figure 7A**, appear critical for maintaining the overall structural stability and functional integrity of PBP2a. However, significant differences in the salt bridge networks were observed between the room-temperature and cryogenic structures. Specifically, the room-temperature XFEL structures exhibit a greater number of salt bridges overall compared to the cryogenic apo PBP2a structure (PDB 1VQQ) (Lim & Strynadka, 2002). In our room-temperature structures, an average number of 91 salt bridges were identified, whereas the cryogenic apo PBP2a structure contained an average of 86 salt bridges (**Table S4**). This observation aligns with a trend reported in a recent study by our group, which also noted an increase in interactions in room-temperature structure of the human NQO1 compared to those observed in cryogenic environments (Grieco *et al*., 2024). The enhanced interaction density observed in the XFEL structures likely reflects the greater conformational flexibility of the protein under near-physiological conditions.

**Figure 7.**
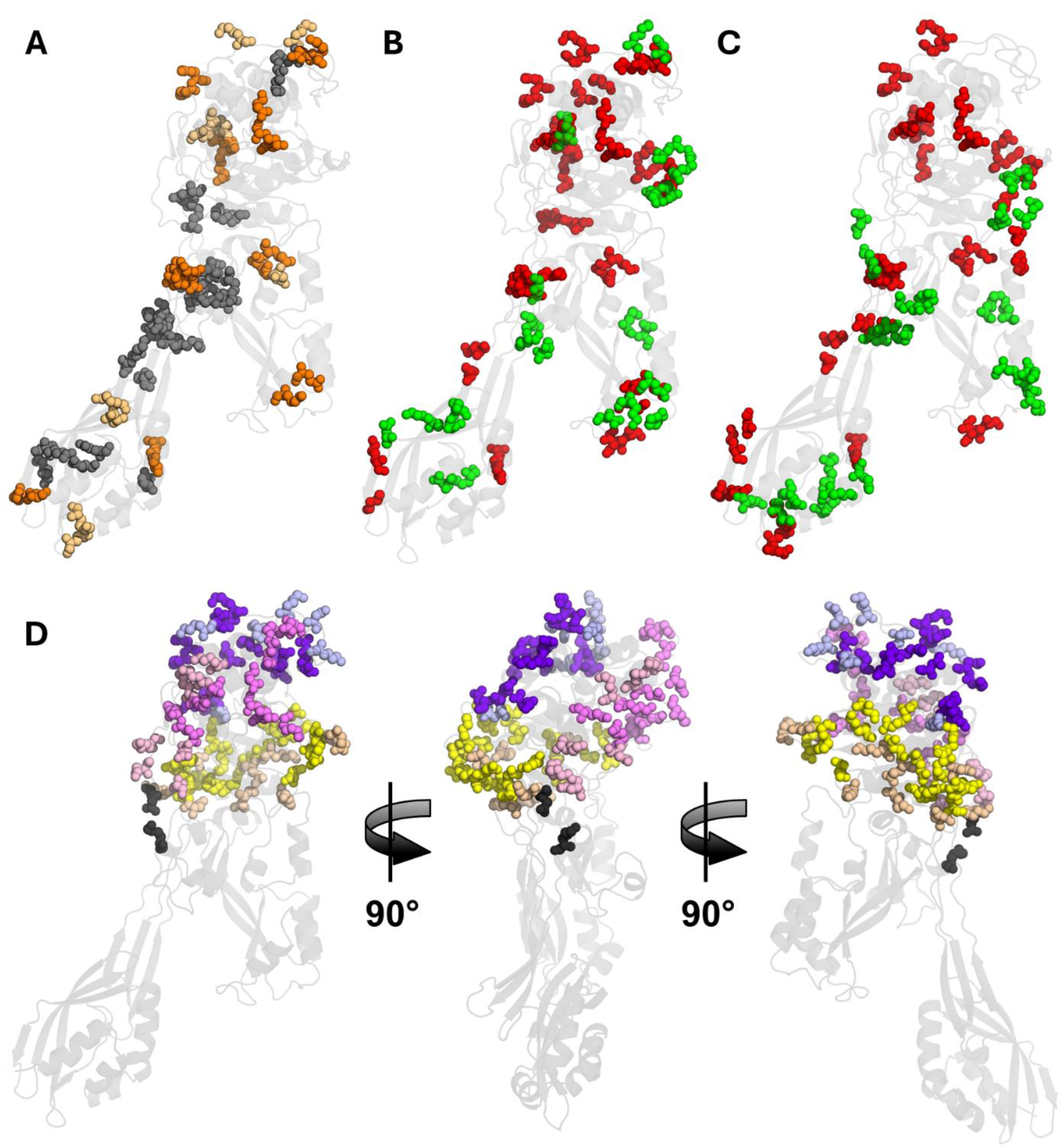
Salt bridge interactions. **A)** The 18 conserved salt bridges shared across all structures are depicted in grey. The 17 novel salt bridges identified exclusively in either chain of the XFEL structures and absent in all chains of the cryogenic apo structure (PDB 1VQQ) (Lim & Strynadka, 2002) are shown in light orange. The 12 salt bridges not observed even in molecular dynamics (MD) simulations (Fishovitz et al., 2014; Mahasenan et al., 2017) are highlighted in orange. **B)** The 24 salt bridges unique to either chain A of the XFEL structures and absent in the PBP2a-ceftaroline complex (PDB 3ZFZ) (Otero et al., 2013) are shown in red, representing those disrupted upon binding. Conversely, the 12 salt bridges absent in the XFEL structures but present in chain A of the PBP2a-ceftaroline complex (PDB 3ZFZ) (Otero et al., 2013) are shown in green, indicating newly formed interactions upon binding. **C)** The 22 salt bridges found in either chain A of the XFEL structures but absent in the PBP2a-PG complex (PDB 3ZG5) (Otero et al., 2013) are shown in red, representing disrupted interactions upon binding. The 13 salt bridges absent in the XFEL structures but present in chain A of the PBP2a-PG complex (PDB 3ZG5) (Otero et al., 2013) are shown in green, representing newly formed interactions upon binding. **D)** Salt bridge interactions found at the back of PBP2a. In all panels, PBP2a is represented as a grey cartoon. The detailed list of all salt bridge interactions represented in this figure is provided in Table S6.

In both chains of the room-temperature XFEL structures, novel salt bridges were identified that are absent in the cryogenic apo PBP2a structure (PDB 1VQQ) (Lim & Strynadka, 2002). These include 17 unique interactions distributed across the PBP2a surface (**Tables S5 and S6,** and **Figure 7A**). Intriguingly, 12 of these novel interactions (**Tables S5 and S6,** and **Figure 7A**) were not even previously identified as one of the 59 most dynamic salt-bridge interactions in molecular dynamics simulations conducted by Mahasenan *and colleagues* (Mahasenan *et al*., 2017). These unique salt bridges may represent transient interactions facilitated by the increased conformational flexibility of PBP2a at room temperature, as captured by XFEL methods. This flexibility contrasts with the more stabilized conformational states observed under cryogenic conditions, where freezing limits the dynamic behavior of the protein and restricts the formation of certain interactions.

In addition to the above differences in the apo structures, we extended our analysis to the room-temperature structures by comparing them to those bound to the antibiotic ceftaroline and peptidoglycan. Specifically, we focused on examining how the salt bridge network evolved upon the binding of these two ligands. Our findings revealed 34 salt bridges that play specific roles in the antibiotic’s interaction with PBP2a (**Table S4**). Of these, 24 interactions are disrupted upon ceftaroline binding (**Tables S5 and S6,** and **Figure 7B**), while 12 new interactions are formed (**Tables S5 and S6,** and **Figure 7B**). These salt bridges are crucial for the allosteric regulation of PBP2a, influencing both ligand binding and the communication between allosteric and catalytic sites. In the case of PG, 22 interactions were disrupted, and 13 interactions were newly formed upon its binding (**Tables S5 and S6,** and **Figure 7C**). Overall, the varied patterns of salt bridge interactions across the analyzed structures underscore the complexity of PBP2a’s functional dynamics and the importance of ligand-induced changes in its regulation.

Notably, specific residues such as Asn146 and Glu150 at the allosteric site have been described to play a central role in facilitating long-range allosteric communication within PBP2a (Fishovitz *et al*., 2014). Mutational studies have shown that disruptions to the salt bridges involving these residues impair the structural connectivity required for transmitting conformational signals from the allosteric to the catalytic site. Structural analysis of the room-temperature XFEL structures revealed that neither Asn146 nor Glu150 engages in direct interactions with immediately adjacent residues (**Table S4**). This observation aligns with their proposed role as "allosteric nodes," positioned to mediate structural communication by facilitating transient interactions or propagating conformational shifts across the protein. In this context, we have also identified two residues that may play a role in signal propagation. Residues Lys290 in the allosteric domain and Asp552 in the transpeptidase domain form a salt bridge that connects these two domains. This salt bridge is present in the room-temperature structures and the structure with peptidoglycan (PDB 3ZG5) but is absent in the structure with ceftaroline (**Table S5**).

## 4. Discussion

### 4.1. PBP2a is a suitable protein for time-resolved experiments at XFELs

PBP2a represents a challenging yet promising protein that we have successfully adapted for TR-SFX studies at XFELs, thanks to extensive troubleshooting and optimization efforts. Although TR-SFX studies on PBP2a have not yet been performed, this work establishes a strong foundation for future investigations into its dynamic mechanisms. The protein’s inherent flexibility, particularly within its active and allosteric sites, makes it an excellent candidate for capturing conformational transitions critical to its enzymatic function through TR experiments. Iterative refinement of expression, purification, and crystallization protocols enabled the production of high-quality samples tailored for serial crystallography studies. Expression strategies yielded highly purified PBP2a at rates exceeding 100 mg per week. Batch crystallization under continuous agitation produced uniform microcrystals with excellent diffraction quality, optimized further for sample delivery using injectors such as the MESH and the DFFN. Initial static SFX experiments demonstrated the feasibility of studying PBP2a at XFELs and highlighted the unique advantages of this technology. Cd^2+^ ions, which stabilize the enzyme’s dimeric form, pose significant challenges due to their increased susceptibility to radiation damage. XFEL’s ultrafast pulses addressed this issue by capturing diffraction data before radiation-induced damage could occur, thereby preserving the structural integrity of Cd^2+^-stabilized crystals. These experiments underscored PBP2a’s potential for TR-SFX studies. Future TR-SFX investigations could employ the mix-and-inject approach to monitor real-time enzymatic events, including ligand binding, allosteric transitions, and active site rearrangements. Capturing short-lived intermediate states would provide critical insights into PBP2a’s mechanism of resistance to β-lactam antibiotics. Such knowledge could inform the design of novel inhibitors targeting transient states, potentially overcoming antibiotic resistance. By demonstrating the feasibility of adapting PBP2a for XFEL-based studies and establishing the groundwork for TR-SFX experiments, this study opens new avenues for exploring the unique allosteric mechanisms of PBP2a. Capturing its conformational transitions in real time would yield a deeper understanding of its enzymatic processes and further validate TR-SFX as a transformative tool for studying dynamic proteins. The foundation laid here represents a significant step toward addressing critical questions about PBP2a’s function and developing therapeutic strategies to combat multidrug-resistant bacterial infections.

### 4.2. Effect of Cd^2+^ ions on PBP2a crystal formation and crystal damage

Cd^2+^ ions play a crucial role in PBP2a dimerization and crystal formation by bridging negatively charged residues, stabilizing specific protein conformations, and promoting crystal lattice formation. Like other divalent cations, Cd^2+^ enhances electrostatic interactions and facilitates dimerization by coordinating with residues such as histidine and cysteine (Ames *et al*., 1998; Linsdell, 2015). Additionally, it influences crystallization by altering protein solubility, supporting specific crystal contacts, and stabilizing the protein structure. However, Cd^2+^ also presents challenges in X-ray crystallography, as its high atomic number increases radiation absorption, leading to structural damage. X-ray exposure can displace Cd²⁺ ions from binding sites, disrupt disulfide bridges, and degrade the crystal lattice, ultimately reducing diffraction quality (Hekstra, 2023). Despite these challenges, Cd²⁺ remains valuable for promoting crystallization, albeit with careful experimental considerations. Ionization of heavy metal ions under X-ray exposure can distort protein structures, complicating diffraction data analysis and reducing resolution.

Traditionally, cryoprotection—cooling crystals to very low temperatures—has been used to reduce radiation damage by minimizing atomic movement and free radical formation. However, this approach is less effective for heavy metals due to their higher atomic numbers, making them more susceptible to radiation damage. Another strategy involves using softer X-rays to mitigate damage, though this typically results in lower-resolution data. A more effective solution is XFELs, which deliver ultra-short, intense X-ray pulses that capture diffraction data before significant radiation damage occurs (Chapman *et al*., 2011; Neutze *et al*., 2000). This technique, known as serial femtosecond crystallography (SFX), enables data collection from a large number of microcrystals in rapid succession, allowing researchers to outrun radiation damage and preserve crystal integrity (Chapman *et al*., 2011). XFEL-based SFX has proven particularly effective for studying delicate metal-rich proteins that are prone to radiation damage, as it allows for the collection of high-quality structural data (Suga *et al*., 2020; Kern *et al*., 2018; Coe & Fromme, 2016; Wiedorn *et al*., 2018; Sauter *et al*., 2020; Hough & Owen, 2021; Kern *et al*., 2015).

In our case, many attempts to solve the structure of PBP2a using synchrotron-based serial crystallography were unsuccessful, likely due to the radiation damage caused by Cd^2+^ ions. These ions, which stabilize the two PBP2a molecules in the crystal lattice, are particularly sensitive to X-ray exposure. The high radiation absorption by these metal ions leads to structural degradation and disruption of the crystal lattice, making it difficult to obtain accurate diffraction data. However, using XFELs, we successfully solved the room-temperature structure of PBP2a, capturing high-quality data before significant radiation-induced damage could occur. Our findings support the hypothesis that the presence of heavy metals, particularly Cd^2+^ ions acting as molecular bridges between protein molecules, is a key factor in the pronounced radiation damage observed under synchrotron X-ray exposure. This highlights the advantages of XFEL-based serial crystallography for studying metal-rich, radiation-sensitive protein systems. While heavy metals can stabilize protein crystals, their propensity to exacerbate radiation damage underscores the importance of alternative techniques, such as XFELs, to obtain high-quality structural data from fragile protein crystals.

### 4.3. Structural Differences in PBP2a: Comparing Cryogenic and Room-Temperature Conditions

An intriguing observation worth highlighting is that, among the two structures reported by Otero *and colleagues* for PBP2a bound to ceftaroline (PDBs 3ZFZ and 3ZG0) (Otero *et al*., 2013), the structure obtained from soaked crystals (PDB 3ZFZ) showed only one molecule of the two PBP2a in the ASU occupied by the antibiotic at the allosteric and catalytic sites. In contrast, the co-crystallized structure (PDB 3ZG0) displayed both molecules of PBP2a occupied by the antibiotic in the allosteric site, with none in the catalytic site. This discrepancy can be attributed to the fact that cryo-crystals, and cryogenic temperatures in general, can induce conformational changes that reflect a limited subset of the possible conformations at room temperature. In this context, a structural comparison between our room temperature structures of PBP2a at XFELs in its apo form and those obtained under cryogenic conditions (PDB 1VQQ) revealed significant conformational differences, particularly in flexible loop regions and the allosteric site. The room-temperature structures exhibit increased mobility in key residues associated with catalytic site accessibility, suggesting a dynamic equilibrium that may be restricted in cryogenic structures. Furthermore, certain salt bridges observed in the cryogenic structures appear weaker or absent in our data, likely due to temperature-dependent stabilization effects. These findings reinforce the idea that cryogenic conditions may obscure physiologically relevant conformational states, emphasizing the importance of room-temperature structural studies for a comprehensive understanding of PBP2a’s functional dynamics. Differences between room-temperature and cryogenic crystallography structures have been experimentally observed in several proteins (Liu *et al*., 2013; Pan *et al*., 2022; Ayan *et al*., 2022; Wolff *et al*., 2020; Srinivas *et al*., 2020; Suno *et al*., 2018; Fenalti *et al*., 2015; Skaist Mehlman *et al*., 2023). In a recent study by our group, we reported on the room temperature structure of human NQO1 (NAD(P)H oxidoreductase 1), a highly dynamic and allosteric enzyme similar to PBP2a, using XFELs (Doppler *et al*., 2023). For the first time, we observed that NQO1’s two active binding sites act cooperatively, displaying highly collective inter-domain and inter-monomer communication and dynamics. This underscores the importance of determining room temperature structures, as the structural and dynamic information often eludes standard macromolecular crystallography at cryogenic conditions, preventing researchers from observing such behavior from a structural perspective.

### 4.4. New salt bridge network proposed for the opening of the catalytic site

In this study, we have identified two residues that may contribute to signal propagation in PBP2a. Lys290 in the allosteric domain and Asp552 in the transpeptidase domain form a salt bridge that connects these two domains. This salt bridge is observed in both the room-temperature structures and the peptidoglycan-bound structure (PDB 3ZG5) (Otero *et al*., 2013) but is absent in the ceftaroline-bound structure (**Table S5**). Building upon previously known interactions, we propose a complex network of salt bridges linking the allosteric site to the catalytic site. This network, located at the "back" of the enzyme (**Figure 7D**) and opposite the salt bridge network described by Otero *et al*. (Otero *et al*., 2013), consists of 27 salt bridges (**Table S6, Figure 7D**) that undergo significant rearrangement upon ligand binding. Ligand binding, whether to ceftaroline or peptidoglycan, appears to induce reorganization of these interactions, involving the formation and disruption of specific salt bridges that span the transpeptidase domain. Notably, the Lys290-Asp552 salt bridge directly links the allosteric and transpeptidase domains. However, this interaction was not among the 59 most favorable salt bridges identified in the MD simulations by Mahasenan et al. (Mahasenan *et al*., 2017), highlighting its potential relevance. This bridge is present in the peptidoglycan-bound structure but absent in the ceftaroline-bound structure.

Antibiotic binding triggers signal propagation from the allosteric site to the catalytic site, accompanied by increased plasticity in the transpeptidase domain, facilitating antibiotic entry. By contrast, peptidoglycan binding lacks this effect. This dynamic reorganization, including conformational changes in the α_2_-α_3_ loop and/or the α_9_ helix, suggests a role for the salt-bridge network in modulating the catalytic site’s opening. Molecular dynamics simulations by Hermoso’s and Shariarh’s groups (Mahasenan *et al*., 2017; Fishovitz *et al*., 2014) support this hypothesis, which we plan to further test through mix-and-inject time-resolved experiments to investigate the network’s dynamic behavior and its role in driving PBP2a’s catalytic activity.

## 5. Conclusions

PBP2a exemplifies a challenging yet promising protein that has been successfully adapted for time-resolved SFX (TR-SFX) studies at XFELs through extensive troubleshooting and optimization. Its intrinsic structural flexibility, particularly within the active and allosteric sites, makes it an ideal candidate for capturing transient conformational changes crucial to its enzymatic function. The room-temperature crystallographic structures presented in this study offer novel insights into PBP2a’s functional dynamics, highlighting differences from traditional cryogenic structures and providing a clearer understanding of its antibiotic resistance mechanisms.

A key aspect of PBP2a’s structural stability and crystallization is the role of metal ions, which influence electrostatic interactions, protein dimerization, and solubility. In PBP2a, Cd^2+^ ions act as molecular bridges, stabilizing the dimeric form and promoting crystal lattice formation. However, these same ions pose significant challenges in crystallographic studies due to their high atomic numbers, which exacerbate radiation damage under conventional synchrotron X-ray sources. The high radiation absorption of Cd^2+^ can lead to structural degradation, ion displacement, and disruption of crystal integrity, making it difficult to obtain accurate diffraction data.

This study underscores the advantages of XFEL-based SFX in overcoming these limitations. By delivering ultra-short, intense X-ray pulses, SFX captures diffraction data before significant radiation-induced damage can occur, preserving the structural integrity of metal-stabilized protein crystals. In the case of PBP2a, XFEL technology was essential in resolving its room-temperature structure, allowing the identification of novel interaction networks that were previously obscured under cryogenic conditions. Notably, a distinct salt bridge network was observed at the back of the enzyme, suggesting an alternative mechanism for catalytic site opening compared to the front-site network described by Otero and colleagues. While previous studies proposed that the salt bridges at the front primarily stabilized peptidoglycan binding, the findings here indicate that the back-site network, involving residues in the long loop above the catalytic site, may play a more direct role in facilitating its opening.

The room-temperature structures also revealed a previously unmodeled region near the active site, further emphasizing the importance of studying proteins under physiological conditions. Structural comparisons with cryogenic datasets demonstrated that cryogenic cooling imposes constraints on protein flexibility, leading to alterations in unit cell dimensions and potentially masking biologically relevant conformational states. The enhanced flexibility captured at room temperature provides a more accurate representation of the enzyme’s natural behavior, highlighting dynamic structural elements that could be crucial for its function.

Additionally, these results establish PBP2a as a viable model for future time-resolved XFEL studies, which could further explore its dynamic transitions and resistance mechanisms. By employing mix-and-inject methodologies, real-time enzymatic events such as ligand binding, allosteric transitions, and active site rearrangements could be visualized at atomic resolution. Understanding these transient states is critical for designing next-generation β-lactam antibiotics capable of overcoming PBP2a-mediated resistance. This study not only advances our knowledge of PBP2a’s functional dynamics but also demonstrates the broader potential of room-temperature crystallography in tackling antibiotic resistance at a structural level.

## Data availability

The structural data for the apo structures of PBP2a generated in this study are available in the Protein Data Bank repository (https://www.rcsb.org/) under accession codes PDB 9I9S (PBP2a at LCLS) and 9I9T (PBP2a at EuXFEL).

## Supplemental Information

The following information is provided as supplemental information: Figures S1-S10, and Tables S1-S6

## Author contributions

J.M.M.-G. conceived and designed the experiments. A.G. and I.Q.-M. produced the protein. A.G., I.Q.-M., and J.M.M.-G. crystallized the protein. A.G., I.Q.-M., S.B., K.D., C.S., H.H., M.V., J.S., F.K., R.d.W., K.K., A.R., J.B., C.K., S.L., P.S., R.S., V.M., L.B.G., R.B., S.M., M.C., J.A.H., J.M.M.-G., participated in the XFEL experiments. A.G., I.Q.-M., and J.M.M.-G., S.B., J.O., S.R., D.d.S., participated in the synchrotrons experiments. M.V., R.d.W., A.R., J.B., C.K., S.L., P.S., R.S., V.M., L.B.G., R.B., S.B., J.O., S.R., D.d.S., contributed to instrument set-up and experimental support and participated as beamline scientists. S.B., A.G. and J.M.M.-G. analyzed diffraction data and solved crystallographic structures. A.G. and J.M.M.-G made the figures of the manuscript. A.G., J.A.H., S.M., M.C., and J.M.M.-G. wrote the manuscript with input from all other co-authors.

## Acknowledgements

JO, SBa and DdS would like to acknowledge members of the ESRF-EMBL Joint Structural Biology Group (JSBG) and other support groups in the commissioning and operation of the new ID29. The results reported here from beamline ID29 were collected from the BAG proposal number MX2427 and MX2539. The results reported here from XFEL beamlines MFX and SPB/SFX were collected from protein crystal screening proposals numbers P10005 and P2967, respectively. Use of the Linac Coherent Light Source (LCLS), SLAC National Accelerator Laboratory, is supported by the U.S. Department of Energy, Office of Science, Office of Basic Energy Sciences under Contract No. DE-AC02-76SF00515. Parts of the sample injector used at LCLS for this research were funded by the National Institute of Health grant P41GM139687. We acknowledge European XFEL in Schenefeld, Germany, for provision of X-ray free-electron laser beamtime at SPB/SFX and would like to thank the staff for their assistance. Sedimentation velocity assays were performed at the Molecular Interactions Facility of the Margarita Salas Center for Biological Research (CIB).

