## Supplemental Information for "Ambient temperature structural studies of Penicillin-binding Protein 2a of *Methicillin-Resistant Staphylococcus aureus* with XFELs and synchrotrons"

#### Abstract

Penicillin-binding protein 2a (PBP2a) is a transpeptidase responsible for the β-lactam resistance in methicillin-resistant *Staphylococcus aureus* (MRSA), posing significant challenges to antibiotic therapy. PBP2a's unique structural features, including its highly flexible active site and allosteric regulation, enable it to maintain catalytic activity even in the presence of β-lactam antibiotics. Despite extensive characterization using cryogenic crystallography, key questions remain about its dynamic properties and conformational changes under near-physiological conditions. Room-temperature crystallography methods, particularly serial femtosecond X-ray crystallography (SFX) at XFELs, provide a powerful approach to capture these dynamics. Unlike cryogenic conditions that can constrain protein flexibility, SFX enables the study of conformational variability and interaction networks closer to physiological states. Here, we present the first room-temperature structures of PBP2a obtained using SFX to uncover insights into the enzyme's flexibility, allosteric communication, and catalytic mechanisms. These findings are built upon optimized large-scale production and crystallization protocols for PBP2a, ensuring high-quality microcrystals suitable for SFX data collection. The room-temperature structures reveal novel interaction patterns, including unique salt bridge networks and dynamic structural elements absent in cryogenic studies. Furthermore, comparative analyses highlight how environmental conditions influence the conformational states of PBP2a, providing new perspectives on its resistance mechanisms. By integrating structural data from EuXFEL and LCLS, this study not only enhances our understanding of PBP2a's functional dynamics but also underscores the value of room-temperature crystallography in studying antibiotic resistance. These insights could guide the design of next-generation β-lactam antibiotics capable of overcoming PBP2a-mediated resistance.

#### Table of contents


| | |
|---|---|
| **Figure S1** | Representative diffraction patterns of PBP2a microcrystals. |
| **Figure S2** | SDS-PAGE and chromatographic analysis of PBP2a purification steps. |
| **Figure S3** | Schematic representation of the large-scale expression and purification workflow for PBP2a. |
| **Figure S4** | Crystallization of PBP2a microcrystals using the batch method with agitation. |
| **Figure S5** | PBP2a dimers observed in the crystals. |
| **Figure S6** | Comparison of unit cell dimensions and crystal packing in room-temperature and cryogenic structures of PBP2a. |
| **Figure S7** | Structural Comparison of apo PBP2a with related cryogenic structures. |
| **Figure S8** | Structural alterations at the allosteric site of PBP2a in room-temperature structure PBP2a-EuXFEL compared to related structures. |
| **Figure S9** | Structural alterations at the catalytic site of PBP2a in room-temperature structure PBP2a-EuXFEL compared to related structures. |
| **Figure S10** | Sedimentation coefficient distribution profiles of PBP2a samples |
| **Table S1** | Interatomic Interactions in Room-Temperature vs. Cryogenic PBP2a Structures |
| **Table S2** | Hydrogen Bonds and Salt Bridges in the Allosteric Site of PBP2a |
| **Table S3** | Hydrogen Bonds and Salt Bridges in the Catalytic Site of PBP2a |
| **Table S4** | Salt Bridge Networks Across PBP2a Structures |
| **Table S5** | Comparison of Salt Bridge Interactions in PBP2a Structures |
| **Table S6** | Conserved and Novel Salt Bridges in PBP2a XFEL and Cryogenic Structures |

#### Supplemental Figures

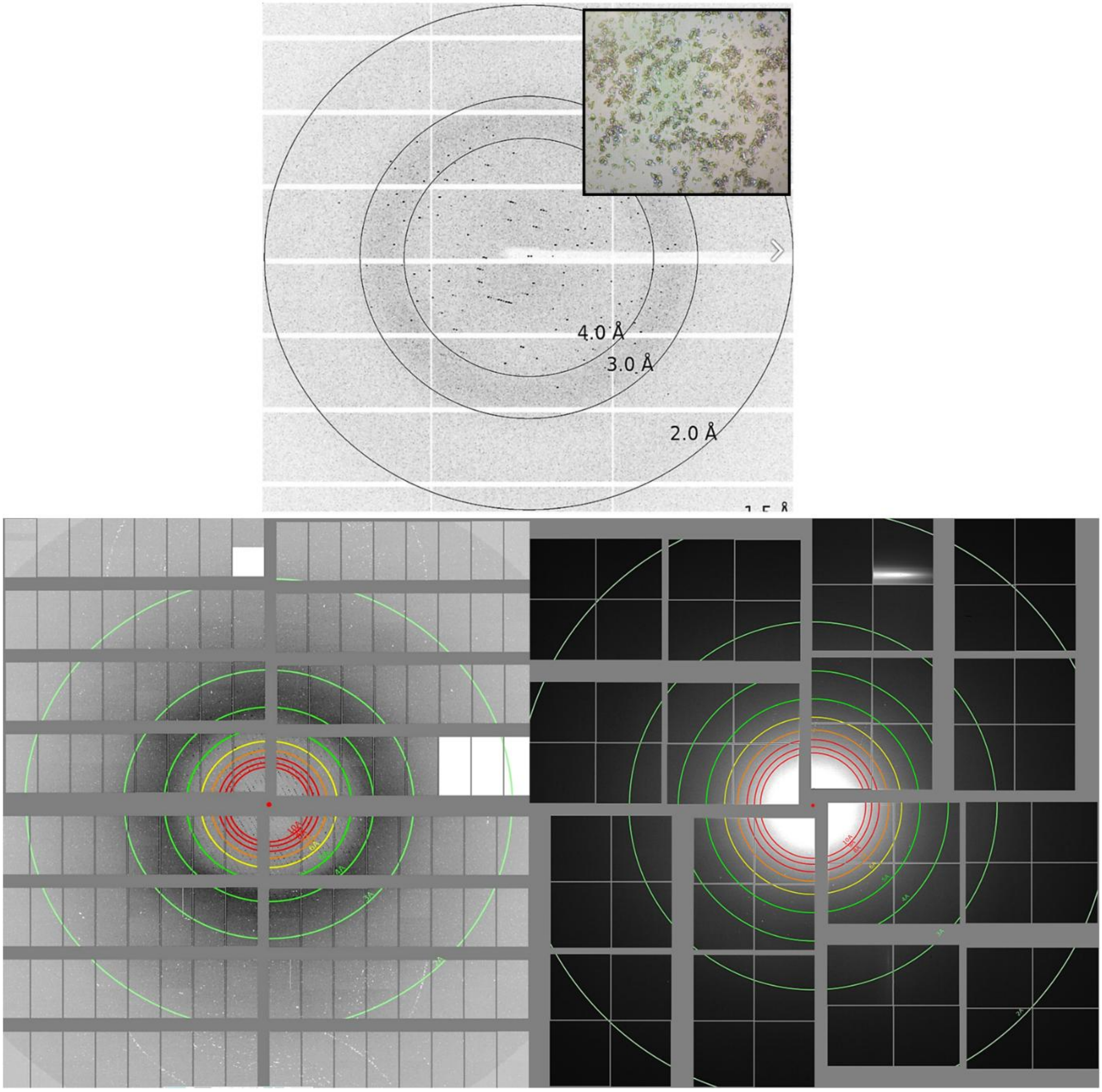


**Figure S1. Representative diffraction patterns of PBP2a microcrystals.** A) Diffraction pattern collected at ID30A from cryo-microcrystals. B) Diffraction pattern collected at MFX. C) Diffraction pattern collected at SPB/SFX.

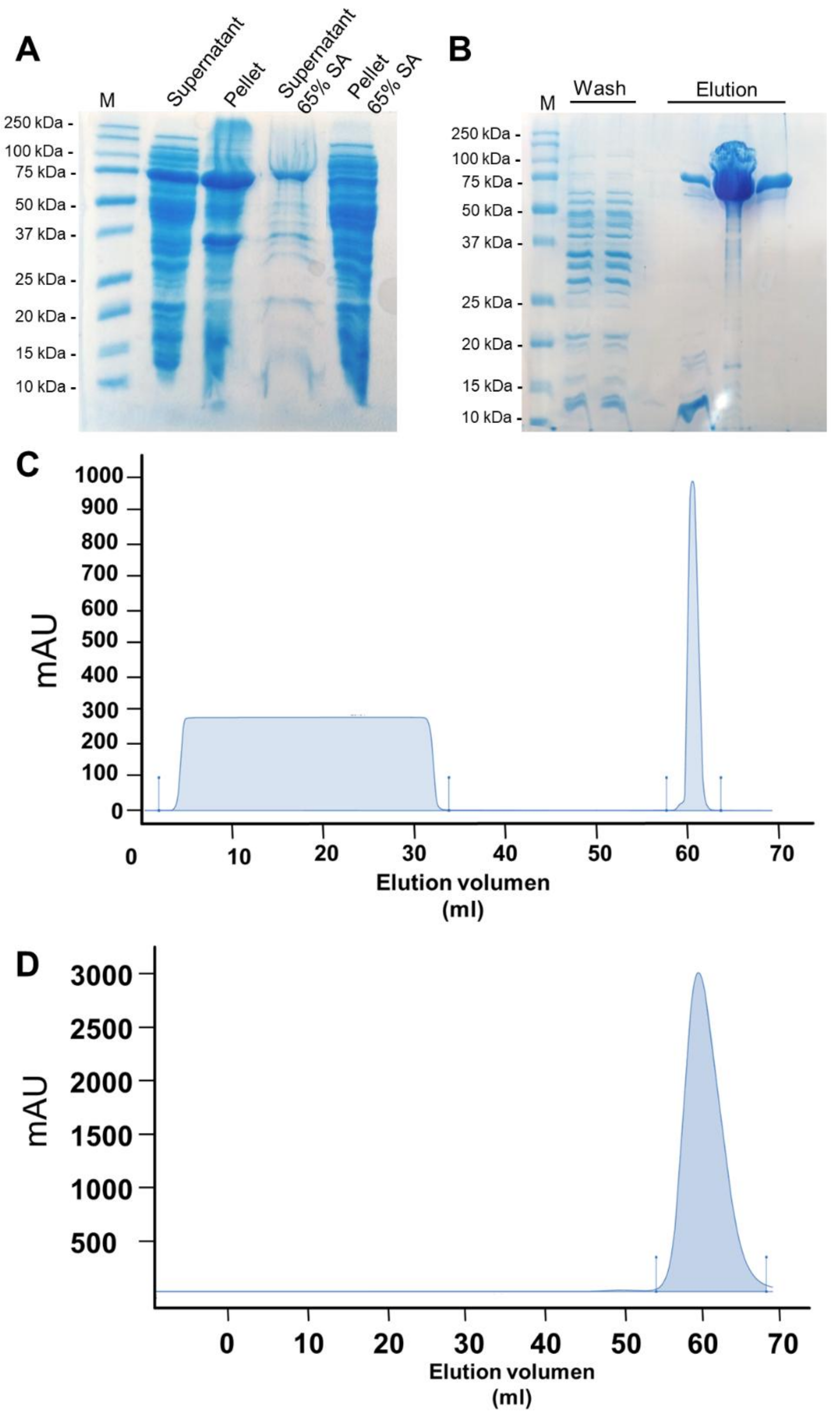


**Figure S2. SDS-PAGE and chromatographic analysis of PBP2a purification steps. A)** SDS-PAGE (12%) showing protein fractions obtained after ammonium sulfate precipitation at 65% saturation, highlighting the enrichment of PBP2a. **B)** SDS-PAGE (12%) of fractions collected following ion exchange chromatography (IEX), demonstrating further purification of PBP2a to near-homogeneity. **C)** Chromatogram of the ion exchange (IEX) purification step, showing the elution profile of PBP2a. **D)** Chromatogram of the size exclusion chromatography (SEC) purification step, confirming the final purification and monodispersity of PBP2a.

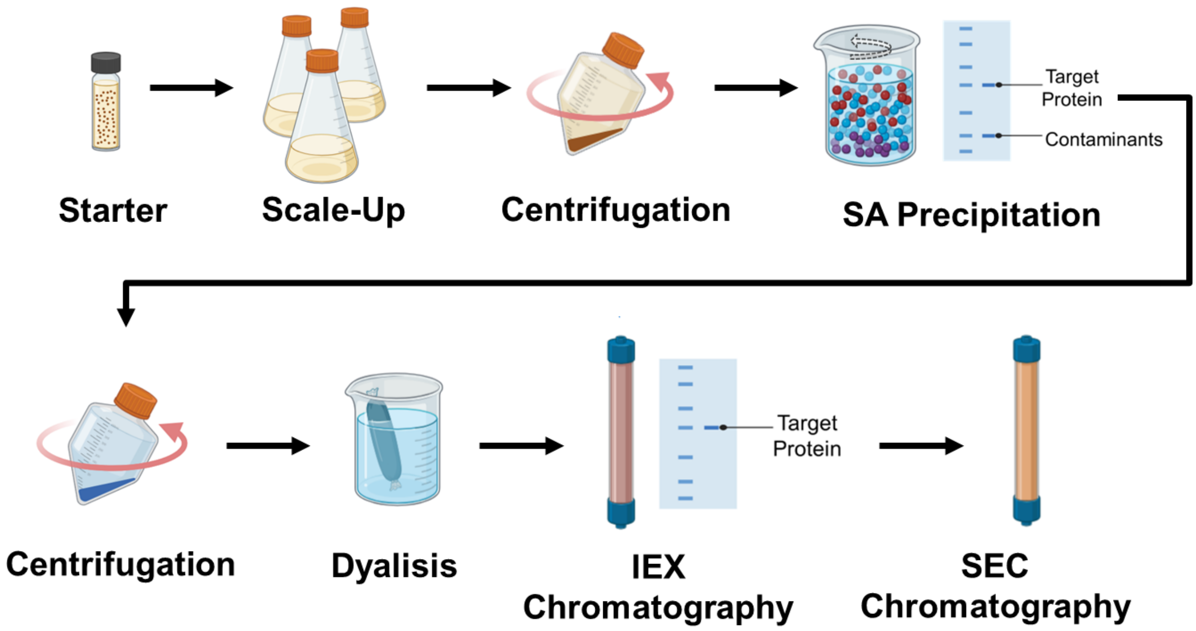


**Figure S3. Schematic representation of the large-scale expression and purification workflow for PBP2a.** The diagram outlines the key steps involved in the overexpression of PBP2a, followed by cell lysis, SA precipitation, protein purification using ionic exchange chromatography (IEX), and size-exclusion chromatography (SEC). This workflow ensures the production of high-purity PBP2a protein suitable for serial crystallography studies.

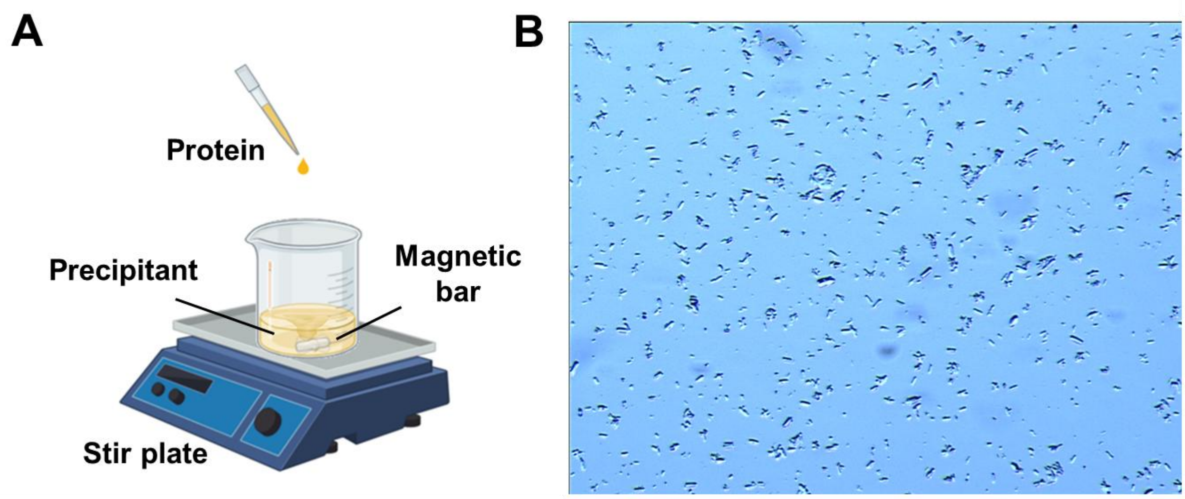


**Figure S4. Crystallization of PBP2a microcrystals using the batch method with agitation. A)** Schematic representation of the batch crystallization setup with continuous agitation. Protein solution is added dropwise to the precipitant solution while stirring at 200 rpm, facilitating the nucleation and growth of microcrystals. **B)** Microcrystals of PBP2a obtained under optimized conditions: 0.1 M HEPES pH 7.0, 25% PEG 1500, 0.88 M NaCl, and 16 mM $CdCl_2$. Crystals reached their final size (~10 µm) within 1 hour at room temperature, demonstrating the efficiency of this method for generating high-quality microcrystals suitable for SSX and SFX experiments.

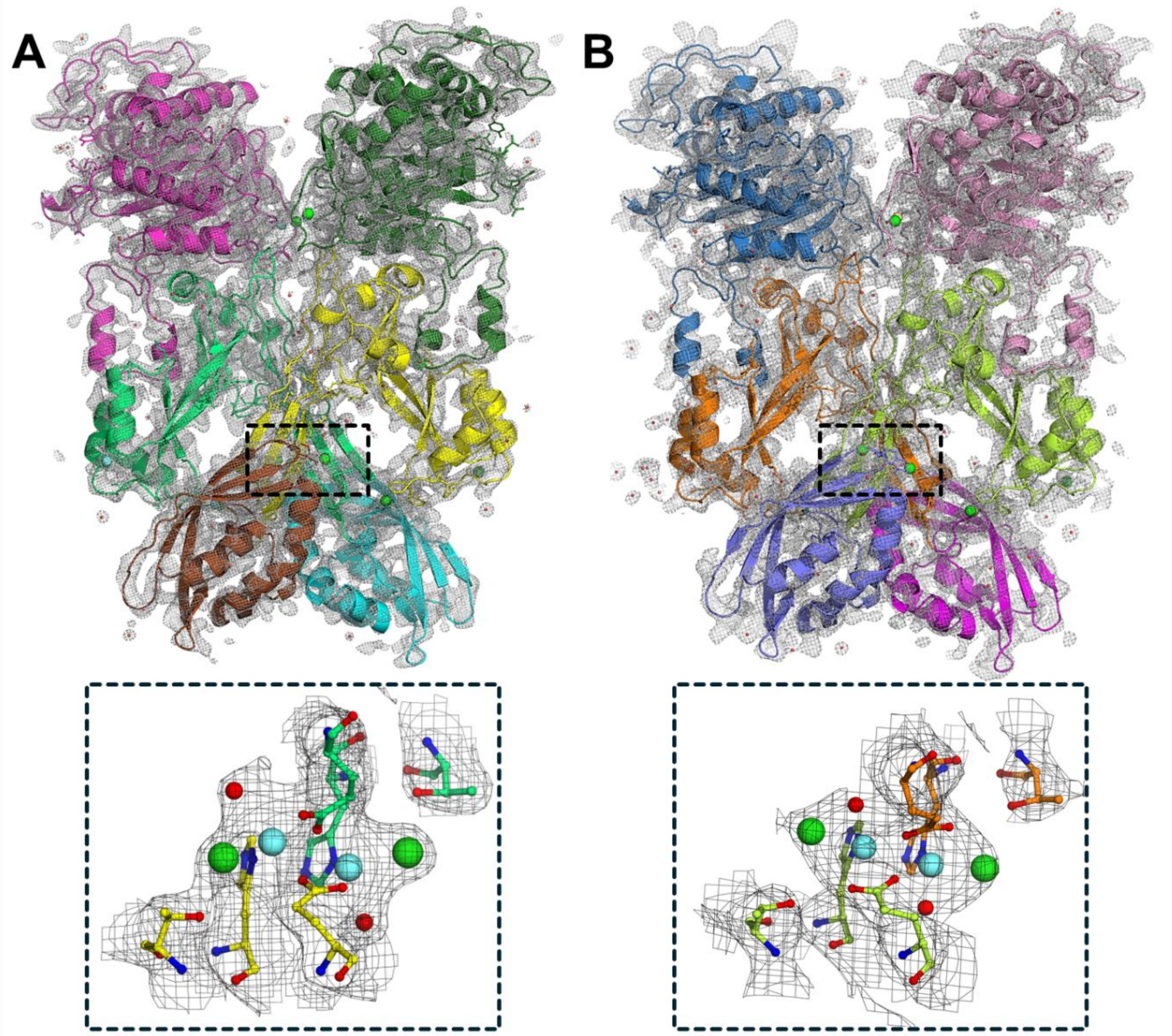


**Figure S5. PBP2a dimers observed in the crystals. A)** Electron 2mF$_o$-DF$_c$ density maps and a cartoon representation of the PBP2a dimer from the room-temperature structure solved at LCLS. The lower panel displays the 2mF$_o$-DF$_c$ density maps surrounding the two $Cd^{2+}$ ions (blue spheres), highlighted with dashed lines in the upper panel. **B)** Electron 2mFo-DFc density maps and a cartoon representation of the PBP2a dimer from the room-temperature structure solved at EuXFEL. The lower panel displays the 2mF$_o$-DF$_c$ density maps around the two $Cd^{2+}$ ions (blue spheres), highlighted with dashed lines in the upper panel.

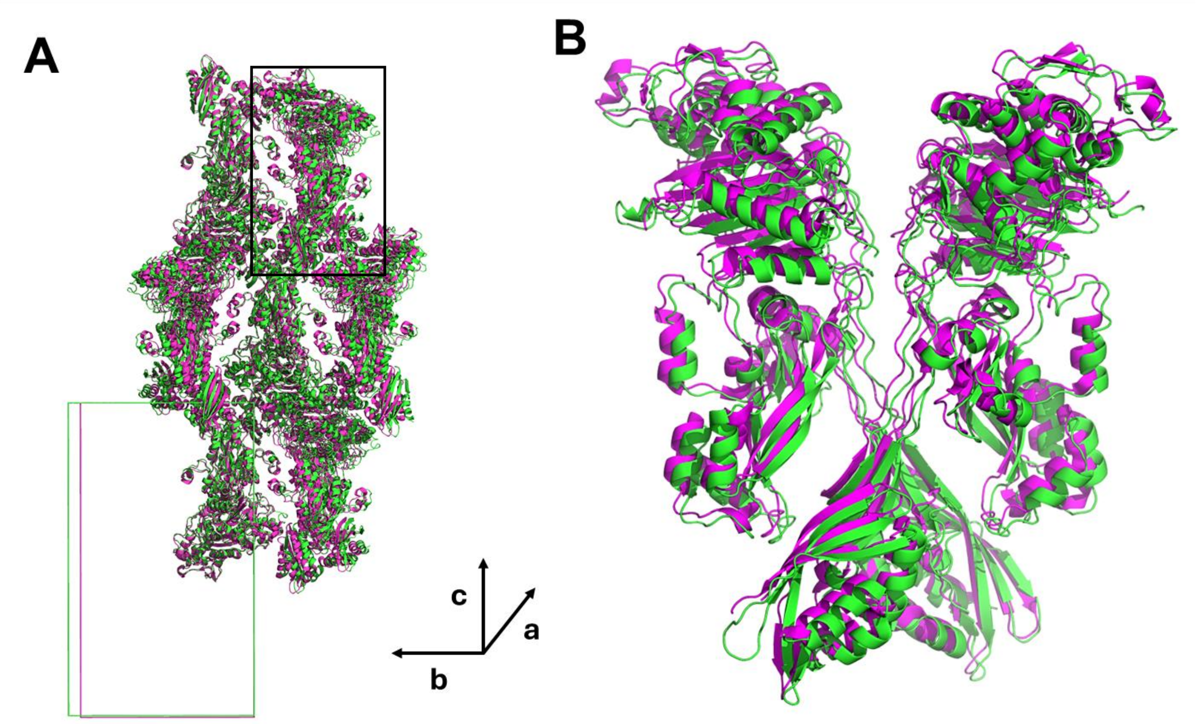


**Figure S6. Comparison of unit cell dimensions and crystal packing in room-temperature and cryogenic structures of PBP2a. A)** Overlay of the crystal packing in the room-temperature structures (green) and the cryogenic structures (magenta), highlighting differences in the b dimension of the unit cell. (**B)** Superimposition of the PBP2a dimer, highlighted with a black box in (A), for the room-temperature structures (green) and the cryogenic structure (PDB 1VQQ) (Lim & Strynadka, 2002) (magenta).

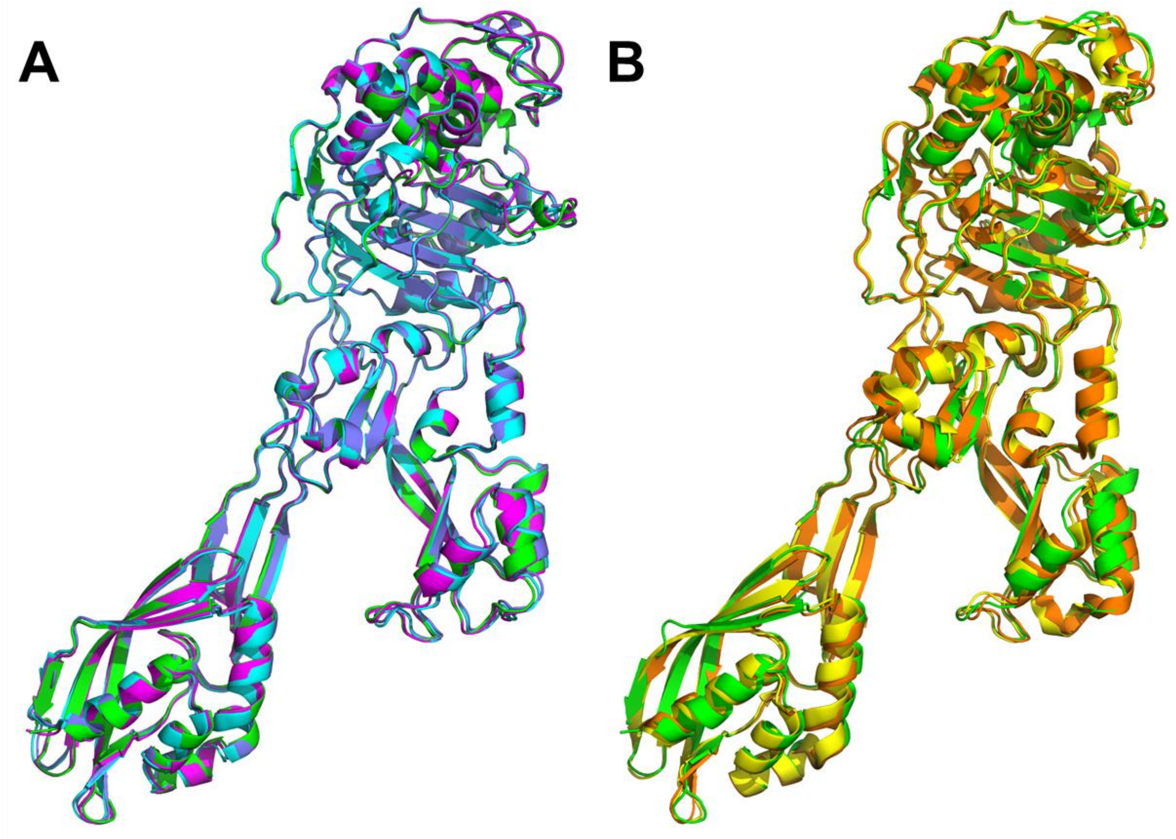


**Figure S7. Structural Comparison of apo PBP2a with related cryogenic structures. A)** Superimposition of all molecules from the room temperature structures reported in this study (chain A of PBP2a-LCLS is shown in green, chain B of PBP2a-LCLS is shown in blue, chain A of PBP2a-EuXFEL is shown in magenta, and chain B of PBP2a-EuXFEL is shown in cyan). **B)** Superimposition of chain A of PBP2a-LCLS (green) to chains A (yellow) and B (orange) of the apo PBP2a structure (PDB 1VQQ) (Lim & Strynadka, 2002).

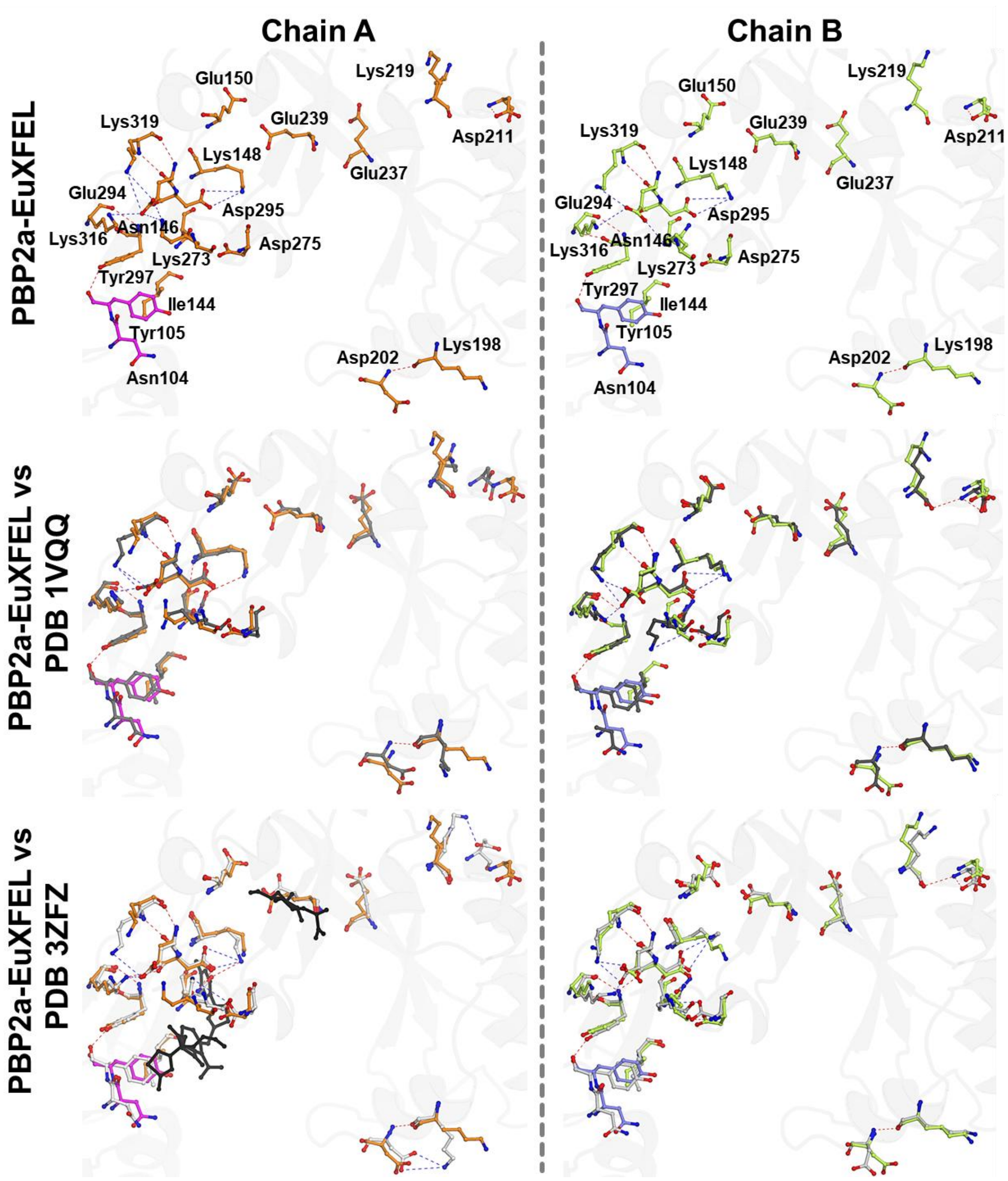


**Figure S8. Structural alterations at the allosteric site of PBP2a in room-temperature structure PBP2a-EuXFEL compared to related structures.** Upper panel depicts all residues forming the allosteric site of PBP2a for chains A and B of the room-temperature PBP2a-EuXFEL structures. Hydrogen bonds within the allosteric site are highlighted in red, while salt bridges are shown in blue. Middle panel depicts the superposition of the allosteric site of PBP2a-EuXFEL with that of the apo structure (PDB 1VQQ) (Lim & Strynadka, 2002). To illustrate the differences in the interaction network with respect to the PBP2a-EuXFEL, only the interactions in (PDB: 1VQQ) (Lim & Strynadka, 2002) are shown using the same color code (hydrogen bonds in red and salt bridges in blue). Lower panel depicts the superposition of the allosteric site of PBP2a-EuXFEL with that of the PBP2a in complex with ceftaroline (PDB 3ZFZ) (Otero *et al.*, 2013). To illustrate the differences in the interaction network with respect to the PBP2a-EuXFEL, only the interactions in (PDB 3ZFZ) (Otero *et al.*, 2013) are shown using the same color code (hydrogen bonds in red and salt bridges in blue). Both ceftaroline and muramic acid ligands are shown as black sticks. For clarity, only residues in the upper panel have been labeled.

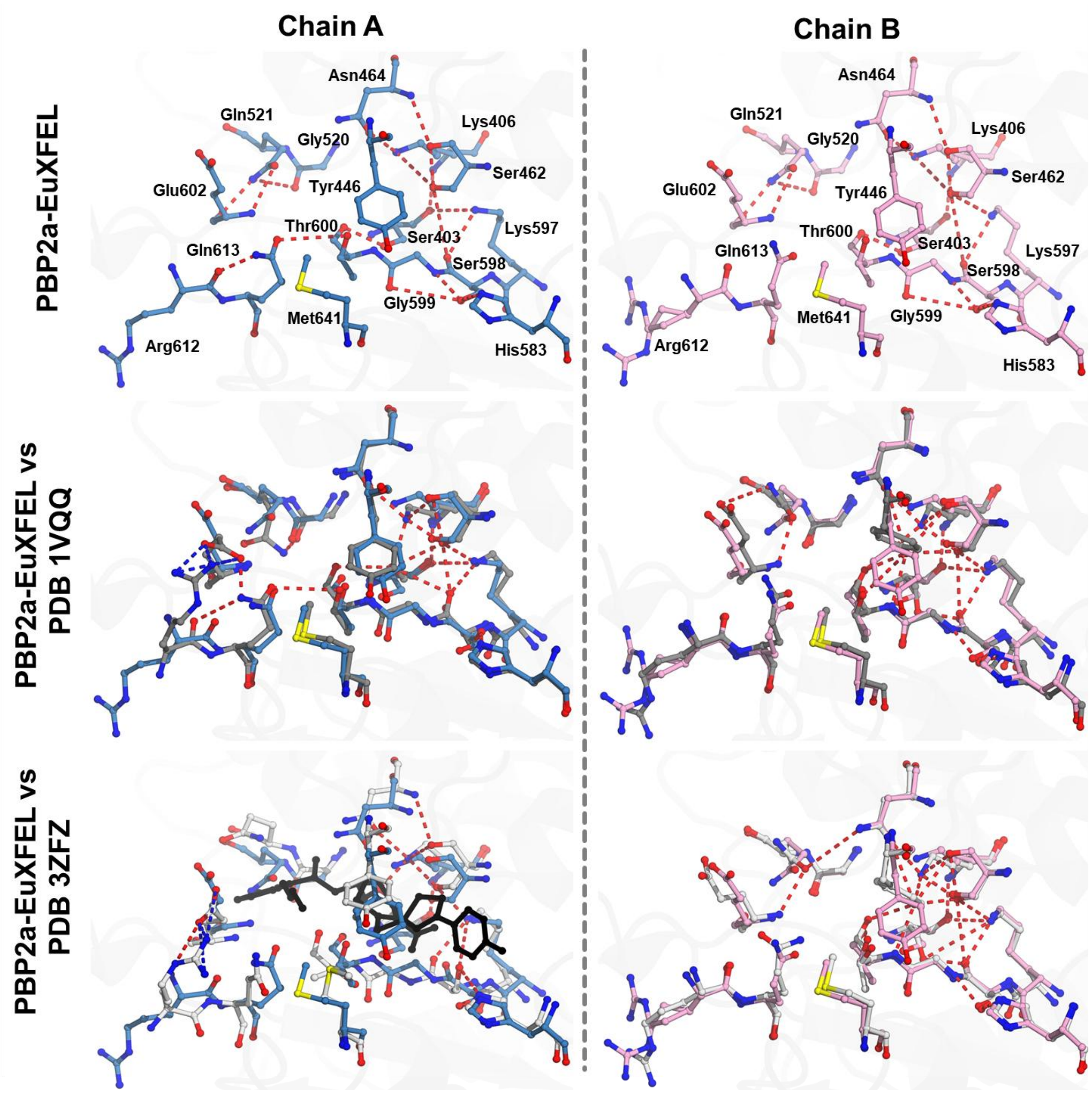


**Figure S9. Structural alterations at the catalytic site of PBP2a in room-temperature structure PBP2a-EuXFEL compared to related structures.** Upper panel depicts all residues forming the catalytic site of PBP2a for chains A and B of the room-temperature PBP2a-EuXFEL structures. Hydrogen bonds within the catalytic site are highlighted in red, while salt bridges are shown in blue. Middle panel depicts the superposition of the catalytic site of PBP2a-EuXFEL with that of the apo structure (PDB 1VQQ) (Lim & Strynadka, 2002). To illustrate the differences in the interaction network with respect to the PBP2a-EuXFEL, only the interactions in (PDB: 1VQQ) (Lim & Strynadka, 2002) are shown using the same color code (hydrogen bonds in red and salt bridges in blue). Lower panel depicts the superposition of the catalytic site of PBP2a-EuXFEL with that of the PBP2a in complex with ceftaroline (PDB 3ZFZ) (Otero *et al.*, 2013). To illustrate the differences in the interaction network with respect to the PBP2a-EuXFEL, only the interactions in (PDB 3ZFZ) (Otero *et al.*, 2013) are shown using the same color code (hydrogen bonds in red and salt bridges in blue). Ceftaroline ligand is shown as black sticks. For clarity, only residues in the upper panels have been labeled.

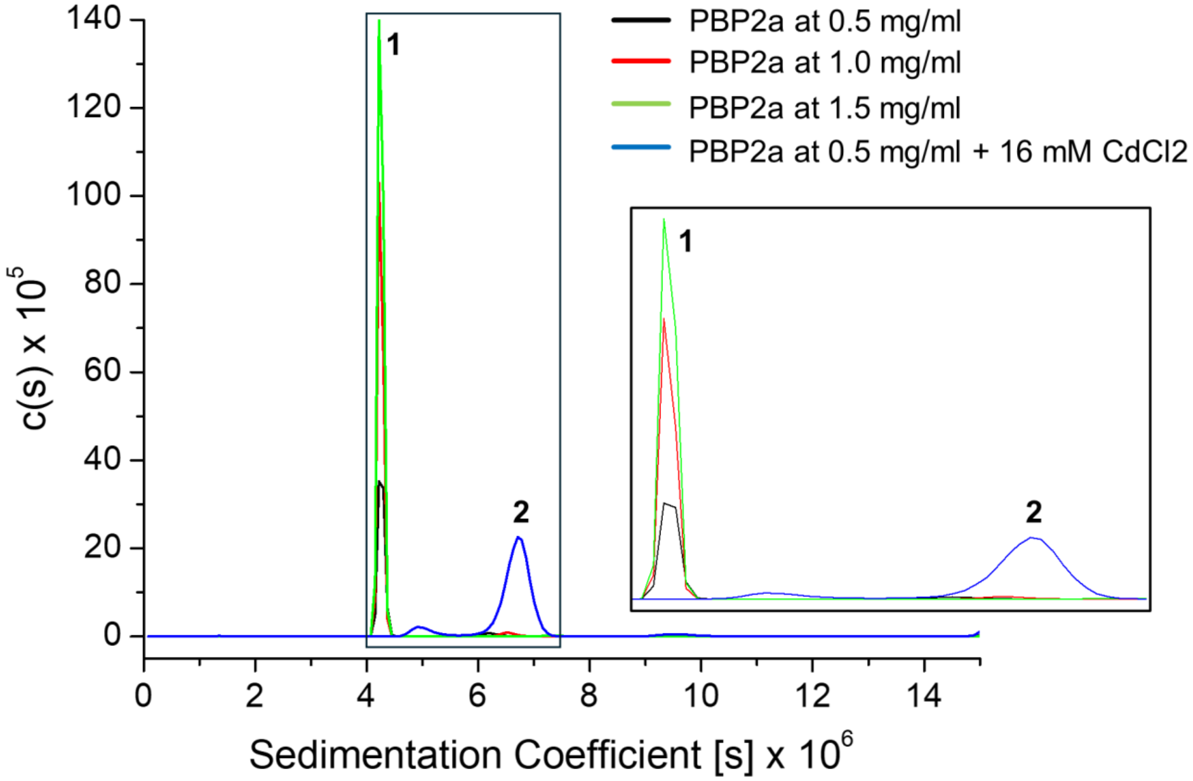


**Figure S10. Sedimentation coefficient distribution profiles of PBP2a samples.** The black, red, and green curves correspond to PBP2a at 0.5 mg/mL, 1.0 mg/mL, and 1.5 mg/mL, respectively. The blue curve represents PBP2a at 0.5 mg/mL in the presence of 16 mM $CdCl_2$. Peaks 1 and 2 indicate different sedimentation species. The inset highlights the main peaks for better visualization.

#### Supplemental Tables

| **Table S1.** Interaction pattern observed in the room temperature structures reported in this study (PBP2a-LCLS and PBP2a-EuXFEL) compared to those solved under cryogenic conditions (PDBs 1VQQ (Lim & Strynadka, 2002) and 3ZFZ (Otero *et al.*, 2013)). | | | | | | | | |
|---|---|---|---|---|---|---|---|---|
| | **PBP2a-LCLS (This study)** | | **PBP2a-EuXFEL (This study)** | | **apo PBP2a (PDB 1VQQ)** | | **PBP2a-Ceftaroline (PDB 3ZFZ)** | |
| | Chain A | Chain B | Chain A | Chain B | Chain A | Chain B | Chain A† | Chain B |
| **Van der Walls interactions** | 1409 | 1346 | 1271 | 1241 | 1786 | 1790 | 1520 | 1504 |
| **Total polar contacts** | 1690 | 1666 | 1545 | 1505 | 2452 | 2422 | 2348 | 2333 |
| Polar contacts | 943 | 943 | 870 | 847 | 1071 | 1064 | 946 | 947 |
| Water mediated polar contacts | 52 | 42 | 54 | 54 | 583 | 594 | 567 | 583 |
| Weak polar contacts | 668 | 660 | 585 | 571 | 637 | 625 | 663 | 646 |
| Water mediated weak polar contacts | 27 | 21 | 36 | 33 | 161 | 139 | 172 | 157 |
| **Total hydrogen bonds** | 1232 | 1215 | 1128 | 1123 | 1865 | 1877 | 1865 | 1888 |
| Hydrogen bonds | 685 | 686 | 626 | 618 | 706 | 694 | 665 | 668 |
| Water mediated hydrogen bonds | 39 | 36 | 44 | 49 | 499 | 513 | 490 | 524 |
| Weak hydrogen bonds | 500 | 482 | 443 | 449 | 524 | 516 | 555 | 547 |
| Water mediated weak hydrogen bonds | 8 | 11 | 15 | 7 | 136 | 154 | 155 | 149 |
| **Salt bridges interactions** | 78 | 83 | 71 | 43 | 80 | 77 | 71 | 89 |
| **Total No. of contacts** | **7331** | **7191** | **6688** | **6540** | **10500** | **10465** | **10017** | **10035** |
| *All the above interactions were calculated using the online version of the ARPEGGIO program* (Jubb *et al.*, 2017). | | | | | | | | |

**Table S2.** Interaction patterns in the allosteric site of PBP2a. Salt bridges (blue) and hydrogen bonds (black) identified in the allosteric site of the apo PBP2a structures from this study are compared to those in the apo structure (PDB 1VQQ) (Lim & Strynadka, 2002) and the ceftaroline-bound complex (PDB 3ZFZ) (Otero *et al.*, 2013).

| | | PBP2a-LCLS (This study) | | PBP2a-EuXFEL (This study) | | Cryo-PBP2a (PDB 1VQQ) | | PBP2a-Ceftaroline (PDB 3ZFZ) | |
|---|---|---|---|---|---|---|---|---|---|
| **Residue/atom 1** | **Residue/atom 2** | **Chain A** | **Chain B** | **Chain A** | **Chain B** | **Chain A** | **Chain B** | **Chain A†** | **Chain B** |
| Asn146 / ND2 | Asp295 / OD2 | -- | -- | -- | -- | -- | 3.2 | 3.3 | 3.2 |
| Asn146 / OD1 | Asp295 / OD2 | -- | -- | -- | -- | 3.6 | -- | -- | -- |
| Lys148 / NZ | Asp295 / OD1 | 3.9 | 3.8 | 4.1 | 4.1 | -- | 4.0 | 4.2 | 3.4 |
| Lys148 / NZ | Asp295 / OD2 | 4.3 | 3.4 | 4.5 | 4.3 | -- | 3.6 | 3.1 | 3.9 |
| Lys148 / N | Asn146 / O | -- | -- | -- | -- | 3.6 | -- | -- | -- |
| Lys148 / NZ | Asn146 / OD1 | -- | -- | -- | -- | 4.0 | -- | 3.7 | -- |
| Lys198 / O | Asp202 / N | 3.2 | 3.0 | 3.2 | 3.2 | 3.4 | 3.1 | 2.9 | 3.5 |
| Lys198 / NZ* | Asp202 / OD1 | -- | -- | -- | -- | -- | -- | 3.3 | -- |
| Lys198 / NZ* | Asp202 / OD2 | -- | -- | -- | -- | -- | -- | 4.3 | -- |
| Lys219 / O | Asp221 / N | 3.8 | 3.9 | -- | -- | -- | 3.7 | -- | 3.7 |
| Lys219 / NZ* | Asp221 / OD1 | -- | -- | -- | -- | -- | -- | 2.9 | -- |
| Asp221 / N | Asp221 / OD1 | 2.9 | 2.9 | 2.7 | 2.7 | -- | -- | -- | 2.8 |
| Asp221 / N | Asp221 / OD2 | -- | -- | -- | -- | -- | 3.7 | -- | -- |
| Lys273 / NZ | Asp275 / OD1 | -- | -- | -- | -- | -- | 4.5 | 4.4 | 3.9 |
| Lys273 / NZ | Asp275 / OD2 | -- | -- | -- | -- | -- | -- | -- | 3.7 |
| Lys273 / NZ* | Glu294 / OE2 | 3.6 | 3.4 | 4.2 | 3.6 | -- | -- | -- | -- |
| Lys273 / NZ | Glu294 / OE1 | -- | -- | 3.1 | -- | -- | -- | -- | -- |
| Asp275 / N | Asp275 / OD1 | 2.7 | 2.7 | 2.7 | 2.7 | 2.7 | 2.7 | -- | 2.9 |
| Glu294 / OE2 | Lys316 / NZ | 3.7 | 4.3 | 3.9 | -- | 2.8 | 2.6 | 2.8 | 3.2 |
| Glu294 / OE1 | Lys316 / NZ | -- | 3.7 | -- | 4.4 | -- | -- | -- | -- |
| Glu294 / O | Lys319 / N | 3.3 | 3.3 | 3.1 | 3.1 | 2.7 | 2.9 | 2.8 | 2.8 |
| Glu294 / N | Lys319 / O | 3.1 | 3.1 | 3.2 | 3.2 | 2.8 | 2.9 | 2.8 | 2.8 |
| Glu294 / OE1 | Lys319 / NZ | 2.9 | 2.9 | 3.8 | 3.8 | 4.3 | 3.3 | -- | 2.9 |
| Glu294 / OE2 | Lys319 / NZ | 3.6 | -- | 4.5 | -- | 3.2 | 3.2 | 3.7 | 3.2 |
| Asp295 / O | Lys316 / NZ | -- | -- | -- | -- | -- | -- | 4.0 | 3.2 |
| Tyr297 / O | Lys316 / N | 3.1 | 3.1 | 3.4 | 3.4 | 2.8 | 2.9 | 2.8 | 2.9 |
| Tyr297 / N | Lys316 / O | 2.9 | 2.9 | 3.2 | 3.2 | 3.0 | 2.9 | 2.9 | 2.7 |
| Tyr297 / OH | Tyr105 / O | -- | -- | 3.4 | 3.4 | 2.7 | 2.9 | 2.7 | 2.9 |

*All shown interactions are in Angstroms.*
*Hydrogen bonds are shown in black and salt bridges are in blue.*
* *Key interactions described in the text are highlighted with asterisk.*
† *Chain A from PDB 3ZF is the only chain in this comparison that has the antibiotic ceftaroline bound to the allosteric and catalytic sites.*

**Table S3.** Interaction patterns in the catalytic site of PBP2a. Salt bridges (blue) and hydrogen bonds (black) identified in the catalytic site of the apo PBP2a structures from this study are compared to those in the apo structure (PDB 1VQQ) (Lim & Strynadka, 2002) and the ceftaroline-bound complex (PDB 3ZFZ) (Otero *et al.*, 2013).

| | | PBP2a-LCLS (This study) | | PBP2a-EuXFEL (This study) | | Cryo-PBP2a (PDB 1VQQ) | | PBP2a-Ceftaroline (PDB 3ZFZ) | |
|---|---|---|---|---|---|---|---|---|---|
| **Residue/atom 1** | **Residue/atom 2** | **Chain A** | **Chain B** | **Chain A** | **Chain B** | **Chain A** | **Chain B** | **Chain A†** | **Chain B** |
| Ser403 / O | Lys406 / N | 3.5 | 3.5 | 3.5 | 3.5 | 3.0 | 3.0 | 3.0 | 3.4 |
| Ser403 / O | Lys597 / NZ | 4.1 | -- | 3.9 | -- | 3.9 | 3.8 | -- | 3.8 |
| Ser403 / OG | Thr600 / O | 2.8 | 2.8 | 2.7 | 2.7 | 3.8 | -- | -- | -- |
| Ser403 / OG | Lys406 / NZ | -- | -- | -- | -- | 2.9 | -- | 2.6 | 4.5 |
| Ser403 / OG | Ser462 / OG | -- | -- | -- | -- | 3.4 | -- | -- | -- |
| Ser403 / OG | Ser598 / O | -- | -- | -- | -- | 3.7 | 2.8 | -- | 2.6 |
| Lys406 / NZ | Tyr446 / OH | -- | -- | -- | -- | -- | -- | -- | 2.9 |
| Lys406 / NZ | Ser462 / O | 3.8 | 3.8 | -- | -- | 2.9 | 3.0 | 3.4 | 3.2 |
| Lys406 / NZ | Ser462 / OG | 3.5 | 3.5 | 3.8 | 3.8 | 2.6 | 2.6 | 2.8 | 2.7 |
| Lys406 / NZ | Asn464 / OD1 | 3.3 | 3.3 | 2.8 | 2.8 | 2.8 | 2.8 | 2.8 | 2.8 |
| Lys406 / NZ | Thr600 / O | -- | -- | -- | -- | -- | 4.2 | -- | -- |
| Tyr446 / OH | Ser462 / O | -- | -- | -- | -- | -- | 3.6 | -- | 3.5 |
| Tyr446 / OH | Ser462 / OG | -- | -- | -- | -- | -- | 3.3 | -- | 2.7 |
| Tyr446 / OH | Asn464 / OD1 | -- | -- | -- | -- | -- | 3.8 | -- | 3.9 |
| Tyr446 / OH | Thr600 / O | -- | -- | -- | -- | -- | 2.8 | -- | 3.0 |
| Tyr446 / OH | Thr600 / N | -- | -- | -- | -- | -- | 3.0 | -- | 3.0 |
| Tyr446 / OH | Thr600 / OG | -- | -- | -- | -- | -- | 3.9 | -- | -- |
| Ser462 / O | Ser462 / OG | 2.6 | 2.6 | 2.6 | 2.6 | 2.9 | -- | -- | 2.8 |
| Ser462 / O | Asn464 / ND2 | 4.1 | -- | -- | -- | -- | -- | -- | -- |
| Ser462 / O | Asn464 / N | -- | -- | 3.1 | 3.1 | -- | -- | 3.3 | -- |
| Ser462 / OG | Lys597 / NZ | 3.6 | 3.6 | -- | 3.4 | 2.9 | 2.8 | 2.9 | 3.0 |
| Ser462 / OG | Ser598 / O | 4.3 | 4.2 | 4.4 | 4.3 | 2.9 | 3.9 | -- | 4.0 |
| Asn464 / ND2 | Asn521 / OE1 | -- | 4.2 | -- | -- | -- | -- | -- | 4.3 |
| Gly520 / O | Asn521 / NE2 | -- | -- | 3.8 | 3.9 | 3.0 | -- | -- | -- |
| Asn521 / NE2 | Glu602 / O | 2.9 | 2.9 | 2.5 | 2.5 | -- | -- | -- | -- |
| Asn521 / NE2 | Glu602 / OE2 | -- | -- | -- | -- | -- | 3.8 | -- | -- |
| Asn521 / OE1 | Glu602 / N | -- | -- | 3.1 | 3.1 | -- | 2.7 | -- | 3.0 |
| His583 / NE2 | Ser598 / OG | 2.6 | 2.9 | 3.2 | 3.2 | 3.1 | 2.9 | 3.4 | 3.0 |
| Lys597 / NZ | Ser598 / O | 2.9 | 2.9 | 3.1 | 3.1 | 2.8 | 3.1 | 4.0 | 2.9 |

| | | | | | | | | | |
|---|---|---|---|---|---|---|---|---|---|
| Ser598 / OG | Ser599 / N | 2.6 | 2.6 | 2.4 | 2.4 | -- | 2.6 | -- | 2.7 |
| Ser598 / OG | Gly599 / O | 4.3 | 4.4 | 4.2 | 4.2 | -- | -- | -- | -- |
| Thr600 / OG1 | Gln613 / OE1 | 3.5 | 3.7 | 4.0 | -- | 3.8 | -- | -- | -- |
| Glu602 / OE1 | Gln613 / NE2 | -- | -- | -- | -- | 2.3 | -- | -- | -- |
| Glu602 / OE1 | Arg612 / NH1 | -- | -- | -- | -- | 4.2 | -- | 4.5 | -- |
| Glu602 / OE1 | Arg612 / NH2 | -- | -- | -- | -- | 3.4 | -- | 2.5 | -- |
| Glu602 / OE1 | Arg612 / NE | -- | -- | -- | -- | -- | -- | 3.3 | -- |
| Glu602 / OE2 | Arg612 / NH1 | -- | -- | -- | -- | 2.8 | -- | -- | -- |
| Glu602 / OE2 | Arg612 / NH2 | -- | -- | -- | -- | 2.8 | -- | 4.0 | -- |
| Arg612 / O | Gln613 / NE2 | 2.8 | -- | 2.5 | -- | 3.7 | -- | -- | -- |

*All shown interactions are in Angstroms.*

*Hydrogen bonds are shown in black and salt bridges are in blue.*

† *Chain A from PDB 3ZF is the only chain in this comparison that has the antibiotic ceftaroline bound to the allosteric and catalytic sites.*

**Table S4.** Salt bridge interaction networks observed across the room-temperature structures of PBP2a reported in this study, as well as the apo structure (PDB 1VQQ) (Lim & Strynadka, 2002) and the PBP2a structures complexed with the antibiotic ceftaroline (PDB 3ZFZ) (Otero *et al.*, 2013) and peptidoglycan (PDB 3ZG5) (Otero *et al.*, 2013).

| | | **PBP2a-LCLS (This study)** | | **PBP2a-EuXFEL (This study)** | | **Cryo-PBP2a (PDB 1VQQ)** | | **PBP2a-Ceftaroline (PDB 3ZFZ)** | **PBP2a-peptidoglycan (PDB 3ZG5)** |
|---|---|---|---|---|---|---|---|---|---|
| **Residue/atom 1** | **Residue/atom 2** | **Chain A** | **Chain B** | **Chain A** | **Chain B** | **Chain A** | **Chain B** | **Chain A†** | **Chain A#** |
| Asp27 / OD1 | Lys93 / NZ | 3.1 | 3.7 | 3.6 | 3.3 | 3.1 | -- | 2.8 | 3.4 |
| Asp27 / OD2 | Lys93 / NZ | 3.1 | 2.7 | 2.6 | 2.9 | 4.7 | -- | 4.7 | -- |
| Asp27 / OD2 | Lys87 / NZ | 3.9 | -- | -- | -- | -- | -- | -- | -- |
| Asp35 / OD1 | Arg83 / NH1 | 2.5 | 2.5 | 2.7 | 2.7 | 3.0 | 2.9 | 3.3 | 3.3 |
| Asp35 / OD2 | Arg83 / NH1 | 4.0 | 4.2 | 4.5 | 4.5 | 4.4 | -- | 4.4 | 4.7 |
| Asp35 / OD1 | Arg83 / NH2 | 2.8 | 2.9 | 2.9 | 2.9 | 2.8 | 2.9 | 3.1 | 2.4 |
| Asp35 / OD2 | Arg83 / NH2 | 4.1 | 4.1 | 4.3 | 4.1 | 4.9 | 4.3 | -- | 4.5 |
| Glu38 / OE1 | Arg83 / NH1 | 4.9 | 5.0 | -- | -- | -- | 4.9 | 5.0 | -- |
| Glu38 / OE1 | Arg83 / NH2 | 2.8 | 2.8 | 2.8 | 2.8 | 3.1 | 2.6 | 3.0 | 3.6 |
| Glu38 / OE2 | Arg83 / NH2 | 4.3 | 4.2 | 4.3 | 4.3 | 3.4 | 3.1 | 3.1 | 4.1 |
| Lys43 / NZ | Glu59 / OE1 | -- | 4.1 | -- | 4.4 | 4.8 | 4.6 | -- | 4.9 |
| Lys43 / NZ | Glu59 / OE2 | -- | 3.4 | 3.8 | 3.8 | 2.8 | 2.6 | 4.5 | 3.3 |
| Asp48 / OD2 | Lys124 / NZ | -- | -- | -- | -- | -- | 4.8 | -- | -- |
| Asp56 / OD2 | Lys138 / NZ | -- | -- | -- | -- | -- | -- | -- | 4.7 |
| Lys43 / NZ | Glu64 / OE2 | -- | -- | -- | -- | 4.3 | 4.7 | -- | 4.3 |
| Glu61 / OE1 | Arg65 / NH2 | 4.7 | 5.0 | 4.6 | 4.8 | -- | -- | -- | -- |
| Glu61 / OE1 | Arg65 / NH1 | 3.2 | 2.8 | 3.0 | 2.7 | 4.2 | 2.8 | 4.1 | 4.2 |
| Glu61 / OE2 | Arg65 / NH1 | 3.5 | 3.9 | 3.6 | 3.6 | 3.8 | 4.2 | 2.6 | 2.7 |
| Glu61 / OE2 | Arg65 / NH2 | -- | -- | -- | -- | -- | -- | 4.8 | 4.8 |
| Arg65 / NH2 | Asp304 / OD2 | -- | 4.6 | 5.0 | 4.7 | -- | -- | -- | -- |
| Asp82 / OD1 | Lys84 / NZ | -- | -- | 5.0 | -- | -- | 4.1 | -- | -- |
| Asp82 / OD2 | Lys84 / NZ | -- | -- | 4.3 | -- | -- | -- | -- | -- |
| Asp82 / OD2 | Lys100 / NZ | -- | -- | -- | -- | 4.8 | -- | 4.1 | -- |
| Lys86 / NZ | Asp96 / OD1 | 3.8 | -- | 4.1 | -- | -- | -- | 4.3 | 2.9 |
| Lys86 / NZ | Asp96 / OD2 | 4.3 | -- | 3.5 | -- | 4.2 | -- | -- | 3.7 |
| Lys92 / NZ | Asp126 / OD2 | -- | -- | -- | -- | -- | -- | -- | 5.0 |
| Arg94 / NH2 | Asp96 / OD1 | 4.0 | 3.4 | 4.2 | 4.4 | 3.2 | 4.4 | -- | 2.9 |
| Arg94 / NH2 | Asp96 / OD2 | 3.0 | 3.3 | 3.5 | 3.6 | 3.6 | 4.9 | 4.6 | 2.5 |
| Arg94 / NH1 | Asp96 / OD1 | -- | 4.1 | -- | 3.1 | 4.9 | 4.8 | -- | 4.4 |
| Arg94 / NH1 | Asp96 / OD2 | -- | 3.3 | -- | 2.8 | -- | 4.1 | -- | 4.6 |

| | | | | | | | | | |
|---|---|---|---|---|---|---|---|---|---|
| Lys102 / NZ | Asp77 / OD1 | -- | -- | -- | 4.1 | -- | -- | 4.9 | -- |
| Lys102 / NZ | Asp77 / OD2 | -- | -- | -- | 5.0 | -- | -- | 3.0 | -- |
| Lys100 / NZ | Asp109 / OD1 | 4.3 | -- | -- | -- | 4.3 | 4.8 | -- | -- |
| Lys100 / NZ | Asp109 / OD2 | 4.5 | -- | -- | 4.8 | -- | 4.0 | 4.9 | 4.6 |
| Lys124 / NZ | Glu119 / OE1 | -- | -- | -- | -- | -- | -- | 3.1 | -- |
| Lys124 / NZ | Glu119 / OE2 | -- | -- | 3.4 | -- | -- | -- | -- | -- |
| Lys124 / NZ | Asp120 / OD1 | -- | -- | 4.0 | -- | 2.6 | -- | 3.8 | -- |
| Lys124 / NZ | Asp120 / OD2 | -- | -- | 2.7 | -- | 3.5 | -- | -- | -- |
| Lys138 / NZ | Asp128 /OD1 | -- | -- | -- | -- | -- | -- | 5.0 | -- |
| Lys138 / NZ | Asp128 /OD2 | -- | -- | -- | -- | -- | -- | 4.9 | -- |
| Lys148 / NZ | Asp295 / OD1 | 3.9 | 3.8 | 4.1 | 4.3 | -- | 4.0 | 4.2 | 4.5 |
| Lys148 / NZ | Asp295 / OD2 | 4.3 | 3.4 | 4.5 | 4.3 | 4.7 | 3.6 | 3.1 | 4.7 |
| Arg151 / NH1 | Asp288 / OD1 | 3.3 | 3.3 | 3.3 | 3.3 | 2.8 | 2.9 | 2.8 | 4.6 |
| Arg151 / NH1 | Asp288 / OD2 | 4.5 | 4.5 | -- | -- | 4.5 | 4.6 | 4.5 | 3.2 |
| Arg151 / NH2 | Asp288 / OD2 | 4.6 | 4.6 | 4.8 | 4.8 | 5.0 | -- | 4.8 | -- |
| Arg151 / NH2 | Asp288 / OD1 | 2.8 | 2.8 | 2.6 | 2.6 | 2.9 | 3.0 | 2.8 | 3.3 |
| Arg151 / NH2 | Glu284 / OE1 | 3.3 | 3.3 | 3.5 | 3.5 | 3.0 | 2.9 | 3.2 | 2.6 |
| Arg151 / NH2 | Glu284 / OE2 | -- | -- | -- | -- | 4.8 | -- | -- | 4.7 |
| Arg151 / NH1 | Glu284 / OE1 | 4.9 | 4.9 | 4.9 | -- | 4.7 | -- | 4.8 | 4.3 |
| Lys153 / NZ | Glu150 / OE1 | -- | -- | -- | -- | -- | -- | 4.8 | -- |
| Lys153 / NZ | Asp323 / OD1 | 4.2 | 4.2 | 4.6 | 4.7 | 4.9 | -- | 4.8 | -- |
| Lys153 / NZ | Asp323 / OD2 | 3.7 | 3.7 | 3.5 | 3.5 | -- | -- | 4.7 | -- |
| Lys153 / NZ | Glu161 / OE1 | -- | 4.8 | 4.9 | 4.5 | 3.4 | 2.9 | -- | 3.8 |
| Lys153 / NZ | Glu161 / OE2 | 4.9 | -- | 4.6 | 4.3 | 4.1 | 3.5 | 4.9 | 4.8 |
| Arg157 / NH1 | Glu356 / OE1 | -- | 3.1 | -- | -- | 2.8 | 2.9 | 2.7 | 2.7 |
| Arg157 / NH2 | Glu356 / OE1 | 4.6 | 2.7 | 4.3 | -- | 3.2 | 2.8 | 3.0 | 2.9 |
| Arg157 / NH1 | Glu356 / OE2 | 3.0 | -- | 3.2 | 5.0 | 4.3 | 2.9 | 4.4 | 4.7 |
| Arg157 / NH2 | Glu356 / OE2 | 3.6 | -- | 2.6 | 4.7 | 4.1 | 3.9 | 3.9 | 4.4 |
| Arg157 / NH1 | Asp667 / OD1 | 4.8 | 4.8 | 4.8 | 4.9 | -- | -- | -- | 4.5 |
| Arg157 / NH1 | Asp667 / OD2 | -- | -- | -- | -- | 4.7 | 4.8 | -- | -- |
| Arg157 / NH2 | Asp667 / OD1 | 3.1 | 3.1 | 3.4 | 3.4 | 4.0 | 4.1 | 3.2 | 2.6 |
| Arg157 / NH2 | Asp667 / OD2 | 4.4 | 4.4 | 4.9 | 4.6 | 2.7 | 2.9 | 4.5 | 3.9 |
| Lys176 / NZ | Asp208 / OD1 | -- | -- | -- | 4.8 | -- | -- | -- | -- |
| Lys176 / NZ | Asp208 / OD2 | 4.3 | 4.7 | -- | 4.8 | -- | 4.2 | -- | -- |
| Lys180 / NZ | Glu194 / OE2 | -- | -- | -- | -- | -- | -- | 4.8 | -- |

| | | | | | | | | | |
|---|---|---|---|---|---|---|---|---|---|
| Lys181 / NZ | Asp182 / OD1 | -- | -- | -- | -- | -- | -- | -- | 3.3 |
| Lys181 / NZ | Asp182 / OD1 | -- | -- | -- | -- | -- | -- | -- | 3.3 |
| Lys184 / NZ | Glu194 / OE1 | -- | -- | -- | -- | -- | -- | -- | 2.7 |
| Lys184 / NZ | Glu194 / OE2 | -- | -- | -- | -- | -- | -- | -- | 4.2 |
| Glu194 / OE1 | Lys198 / NZ | -- | 4.7 | -- | -- | -- | -- | -- | 4.5 |
| Glu194 / OE2 | Lys198 / NZ | -- | 4.5 | 4.8 | 4.2 | -- | -- | -- | 3.6 |
| Asp195 / OD1 | Lys198 / NZ | 4.5 | 4.2 | 4.5 | -- | -- | -- | -- | 4.3 |
| Asp195 / OD2 | Lys198 / NZ | 4.1 | 4.1 | -- | -- | 4.0 | -- | -- | -- |
| Lys198 / NZ | Asp202 / OD1 | -- | -- | -- | -- | -- | -- | 3.3 | -- |
| Lys198 / NZ | Asp202 / OD2 | -- | -- | -- | -- | -- | -- | 4.3 | -- |
| Asp226 / OD2 | Lys230 / NZ | -- | 4.8 | -- | -- | -- | 4.8 | -- | -- |
| Lys219 / NZ | Asp221 / OD1 | -- | -- | -- | -- | -- | -- | 2.9 | 4.2 |
| Lys219 / NZ | Asp221 / OD2 | -- | -- | -- | -- | -- | -- | 4.5 | 2.4 |
| Arg241 / NH2 | Glu284 / OE1 | 3.0 | 3.0 | 3.0 | 3.0 | 3.3 | 3.1 | 3.0 | 3.7 |
| Arg241 / NH2 | Glu284 / OE2 | 3.2 | 3.2 | 3.4 | 3.4 | 3.3 | 3.3 | 3.5 | 3.4 |
| Arg241 / NH1 | Glu284 / OE1 | -- | -- | 5.0 | 5.0 | -- | 5.5 | 5.0 | -- |
| Arg241 / NH1 | Glu284 / OE2 | 4.6 | 4.6 | 4.8 | 4.8 | 4.7 | 4.7 | 4.9 | 4.9 |
| Lys247 / NZ | Glu246 / OE1 | -- | -- | 4.8 | -- | -- | -- | -- | -- |
| Lys247 / NZ | Asp367 / OD1 | -- | -- | 4.5 | -- | -- | | 4.9 | -- |
| Lys247 / NZ | Asp367 / OD2 | -- | -- | 4.1 | -- | -- | -- | -- | 4.6 |
| Glu262 / OE2 | Lys265 / NZ | -- | -- | -- | -- | -- | -- | -- | 4.5 |
| Glu262 / OE1 | Lys265 / NZ | -- | -- | -- | -- | -- | -- | -- | 4.8 |
| Glu263 / OE1 | Lys280 /NZ | 3.0 | 3.0 | 3.1 | 3.1 | 2.7 | 2.8 | 3.7 | 2.2 |
| Glu263 / OE2 | Lys280 /NZ | 2.7 | 2.7 | 2.6 | 2.6 | 2.7 | 3.1 | 2.6 | 2.6 |
| Glu268 / OE1 | Lys285 / NZ | 4.2 | 4.8 | 4.3 | 4.2 | 4.6 | -- | -- | 4.6 |
| Glu268 / OE2 | Lys285 / NZ | -- | -- | -- | -- | 4.9 | -- | -- | -- |
| Glu268 / OE2 | Lys267 / NZ | -- | -- | 4.9 | -- | -- | -- | -- | -- |
| Glu268 / OE1 | Lys267 / NZ | -- | -- | -- | -- | -- | -- | -- | 4.8 |
| Lys273 / NZ | Asp275 / OD1 | -- | -- | -- | 4.8 | -- | 4.5 | 4.4 | 4.5 |
| Lys273 / NZ | Asp275 / OD2 | -- | -- | -- | 4.6 | -- | -- | -- | 4.5 |
| Lys273 / NZ | Glu294 / OE1 | 4.9 | -- | 3.1 | -- | 4.6 | -- | 4.7 | -- |
| Lys273 / NZ | Glu294 / OE2 | 3.6 | 3.4 | 4.2 | 3.6 | -- | -- | -- | -- |
| Lys273 / NZ | Asp295 / OD2 | -- | -- | -- | -- | -- | -- | -- | 4.8 |
| Asp288 / OD1 | Lys289 / NZ | -- | -- | -- | -- | -- | -- | -- | 4.6 |

| | | | | | | | | | |
|---|---|---|---|---|---|---|---|---|---|
| Asp288 / OD2 | Lys289 / NZ | -- | -- | 4.9 | 4.9 | -- | -- | -- | -- |
| Lys290 / NZ | Asp552 / OD1 | -- | 5.0 | -- | -- | -- | 4.3 | -- | -- |
| Lys290 / NZ | Asp552 / OD2 | -- | -- | -- | 4.6 | -- | -- | -- | 3.6 |
| Glu294 / OE1 | Lys316 / NZ | -- | 3.7 | -- | 4.4 | 4.7 | 4.7 | 4.8 | 4.6 |
| Glu294 / OE2 | Lys316 / NZ | 3.7 | 4.3 | 3.9 | -- | 2.8 | 2.6 | 2.8 | -- |
| Arg298 / NH1 | Glu315 / OE1 | 4.2 | -- | 3.7 | -- | 2.7 | 5.0 | 4.8 | 4.5 |
| Arg298 / NH1 | Glu315 / OE2 | -- | 4.5 | 5.0 | -- | 4.7 | -- | 2.8 | 2.7 |
| Arg298 / NH2 | Glu315 / OE1 | -- | -- | -- | 4.5 | 4.9 | 3.7 | -- | -- |
| Arg298 / NH2 | Glu315 / OE2 | -- | -- | -- | -- | -- | 4.8 | 5.0 | 3.7 |
| Glu315 / OE1 | Lys317 / NZ | 4.8 | 3.7 | -- | 3.4 | -- | 4.9 | -- | -- |
| Lys318 / NZ | Asp320 / OD1 | 4.6 | 3.3 | 4.7 | 3.6 | 4.0 | 4.4 | 3.2 | 4.3 |
| Lys318 / NZ | Asp320 / OD2 | 4.7 | 3.2 | 4.5 | 3.5 | 3.4 | 3.4 | 2.9 | 3.2 |
| Lys319 / NZ | Glu294 / OE1 | 2.9 | 3.9 | 3.8 | 3.8 | 4.3 | 3.3 | 4.6 | 2.7 |
| Lys319 / NZ | Glu294 / OE2 | 3.6 | 5.0 | 4.5 | -- | 3.2 | 3.2 | 3.7 | 4.5 |
| Asp329 / OD1 | Lys331 / NZ | 4.5 | 4.5 | 4.2 | 4.2 | 4.0 | 3.3 | 3.6 | 3.4 |
| Asp329 / OD2 | Lys331 / NZ | 3.9 | 3.9 | 3.9 | 3.9 | 4.2 | 2.8 | 3.1 | 2.9 |
| Asp343 / OD1 | Lys639 / NZ | 4.6 | 3.3 | 5.0 | 2.5 | 4.3 | -- | 3.8 | -- |
| Asp343 / OD2 | Lys639 / NZ | -- | -- | -- | 4.2 | -- | -- | -- | -- |
| Lys382 / NZ | Glu378 / OE1 | -- | -- | -- | -- | -- | 4.5 | -- | -- |
| Lys382 / NZ | Glu378 / OE2 | -- | 4.7 | -- | -- | -- | 3.1 | -- | -- |
| Lys382 / NZ | Glu385 / OE2 | -- | -- | -- | -- | -- | 4.8 | -- | -- |
| Lys382 / NZ | Asp386 / OD2 | 3.9 | -- | 4.6 | -- | 4.5 | -- | 4.4 | -- |
| Lys387 / NZ | Asp635 / OD2 | -- | -- | -- | -- | -- | -- | -- | 3.9 |
| Glu389 / OE1 | Lys634 / NZ | -- | -- | -- | -- | 4.7 | 4.8 | -- | 4.8 |
| Asp420 / OD1 | Lys 422 / NZ | 5.0 | -- | -- | -- | -- | -- | -- | -- |
| Lys422 / NZ | Asp421 / OD1 | 3.3 | 4.4 | 3.6 | 4.4 | -- | 3.3 | -- | -- |
| Lys422 / NZ | Asp421 / OD2 | 4.0 | 3.8 | 3.4 | 3.4 | 4.8 | 4.4 | -- | -- |
| Lys426 / NZ | Asp428 / OD1 | -- | -- | -- | -- | 3.4 | -- | 4.3 | -- |
| Lys426 / NZ | Asp428 / OD1 | -- | -- | -- | -- | 3.9 | -- | 4.7 | -- |
| Lys434 / NZ | Asp435 / OD1 | -- | -- | 3.3 | 3.2 | -- | -- | -- | -- |
| Lys434 / NZ | Asp435 / OD2 | 3.9 | 3.9 | -- | -- | -- | 4.6 | -- | 4.8 |
| Lys434 / NZ | Glu511 / OE1 | 2.3 | 3.3 | 2.7 | 4.0 | 2.7 | 2.8 | -- | 4.6 |
| Lys434 / NZ | Glu511 / OE2 | 3.9 | 2.8 | 3.9 | 2.7 | 3.4 | 3.4 | 3.3 | 3.8 |
| Asp435 / OD1 | Lys436 / NZ | -- | -- | -- | -- | -- | -- | -- | 4.5 |

| | | | | | | | | | |
|---|---|---|---|---|---|---|---|---|---|
| Asp435 / OD2 | Lys436 / NZ | -- | -- | 4.6 | -- | -- | | -- | -- |
| Arg445 / NH1 | Asp463 / OD1 | -- | -- | 4.6 | -- | 2.7 | 2.9 | 4.1 | 4.6 |
| Arg445 / NH1 | Asp463 / OD2 | 3.2 | 3.2 | 2.9 | 2.9 | 4.2 | 4.2 | 2.6 | 2.9 |
| Arg445 / NH2 | Asp463 / OD2 | -- | -- | -- | -- | -- | -- | 4.9 | -- |
| Arg445 / NH2 | Asp463 / OD1 | -- | -- | -- | -- | 4.9 | | -- | -- |
| Lys456 / NZ | Asp573 / OD2 | 4.1 | 3.2 | 4.9 | 3.3 | -- | 2.8 | -- | 4.2 |
| Glu480 / OE1 | Lys484 / NZ | 4.4 | 2.9 | 4.3 | 2.9 | -- | 4.7 | -- | -- |
| Glu480 / OE2 | Lys484 / NZ | 3.8 | 4.9 | 4.8 | 4.9 | 3.5 | -- | -- | -- |
| Lys506 / NZ | Asp509 / OD1 | -- | -- | 2.8 | -- | -- | -- | -- | -- |
| Lys506 / NZ | Asp509 / OD2 | -- | -- | 4.9 | -- | -- | -- | -- | -- |
| Lys506 / NZ | Glu523 / OE2 | -- | -- | 5.0 | -- | -- | -- | -- | -- |
| Glu523 / OE1 | Lys604 / NZ | -- | 3.7 | -- | 3.7 | -- | 2.6 | -- | -- |
| Glu523 / OE2 | Lys604 / NZ | -- | -- | -- | -- | -- | 4.7 | -- | -- |
| Lys559 / NZ | Glu490 / OE1 | 3.3 | 3.3 | 4.3 | -- | 2.4 | 2.4 | 2.8 | 3.6 |
| Lys559 / NZ | Glu490 / OE2 | 3.2 | 3.5 | 4.3 | 3.6 | 3.6 | 3.8 | 3.1 | 3.7 |
| Glu539 / OE1 | Lys622 / NZ | 3.3 | 4.2 | 3.0 | 4.9 | -- | -- | -- | -- |
| Glu539 / OE2 | Lys622 / NZ | 4.2 | 3.4 | 2.8 | 4.2 | -- | -- | -- | -- |
| Lys565 / NZ | Glu566 / OE1 | -- | 3.7 | -- | 4.8 | -- | -- | -- | -- |
| Lys581 / NZ | Glu460 / OE1 | 2.7 | -- | 3.0 | 3.0 | 3.4 | -- | 3.6 | 3.4 |
| Lys581 / NZ | Glu460 / OE2 | 3.0 | -- | 3.3 | 3.3 | 4.0 | -- | 3.9 | 4.4 |
| Lys584 / NZ | Glu585 / OE1 | -- | -- | -- | -- | 4.5 | -- | -- | -- |
| Asp586 / OD1 | Lys647 / NZ | 3.4 | 3.0 | 3.5 | 2.6 | 2.7 | 4.2 | -- | -- |
| Arg589 / NH1 | Asp654 / OD1 | -- | -- | -- | 2.4 | -- | 5.0 | -- | -- |
| Arg589 / NH1 | Asp654 / OD2 | 4.5 | 4.4 | 4.9 | 3.4 | -- | 4.9 | 4.9 | -- |
| Arg589 / NH2 | Asp654 / OD1 | 4.5 | 4.5 | 4.4 | 3.5 | 3.8 | 3.8 | 4.1 | 3.9 |
| Arg589 / NH2 | Asp654 / OD2 | 2.7 | 2.7 | 3.0 | 3.0 | 3.1 | 3.0 | 3.0 | 3.1 |
| Arg612 / NH1 | Asp635 / OD1 | 4.8 | 4.1 | 4.7 | -- | -- | 4.9 | -- | 4.4 |
| Arg612 / NH1 | Asp635 / OD2 | 4.9 | 4.6 | 4.8 | 4.9 | -- | 4.8 | -- | -- |
| Arg612 / NH2 | Asp635 / OD1 | 3.3 | 3.6 | 3.6 | 4.3 | -- | 3.5 | -- | 3.1 |
| Arg612 / NH2 | Asp635 / OD2 | 3.6 | 3.0 | 3.2 | 3.1 | -- | 2.7 | -- | 3.1 |
| Arg612 / NH1 | Glu602 / OE1 | -- | -- | -- | -- | 4.2 | -- | 4.5 | -- |
| Arg612 / NH1 | Glu602 / OE2 | -- | -- | -- | -- | 2.8 | -- | -- | -- |
| Arg612 / NH2 | Glu602 / OE1 | -- | -- | -- | -- | 3.4 | -- | 2.5 | -- |
| Arg612 / NH2 | Glu602 / OE2 | -- | -- | -- | -- | 2.8 | -- | 4.0 | -- |
| Asp621 / OD1 | Lys663 / NZ | -- | -- | 4.4 | -- | -- | -- | -- | 4.9 |

| | | | | | | | | | |
|---|---|---|---|---|---|---|---|---|---|
| Asp621 / OD2 | Lys663 / NZ | 3.8 | -- | 3.1 | -- | -- | -- | -- | 3.7 |
| Lys622 / NZ | Asp623 / OD1 | -- | -- | -- | -- | -- | -- | 4.8 | -- |
| Asp623 / OD2 | Lys663 / NZ | -- | -- | 4.9 | -- | -- | -- | -- | -- |
| Lys634 / NZ | Asp635 / OD2 | -- | -- | -- | -- | -- | -- | 3.3 | -- |
| Lys662 / NZ | Glu658 / OE1 | -- | 4.1 | -- | -- | 4.1 | 2.5 | -- | 3.2 |
| Lys662 / NZ | Glu658 / OE2 | -- | -- | 4.9 | 4.9 | -- | 4.3 | 4.1 | -- |
| Lys662 / NZ | Glu668 / OE1 | 4.0 | 4.1 | 3.9 | 3.9 | 3.0 | 3.3 | 3.6 | 3.8 |
| Lys662 / NZ | Glu668 / OE2 | 4.1 | 2.8 | 4.1 | 2.7 | 4.2 | 3.2 | 3.4 | 4.0 |
| Glu668 / OE1 | Lys331 / NZ | 3.7 | -- | 4.7 | -- | 4.3 | -- | -- | 4.2 |
| Glu668 / OE2 | Lys331 / NZ | -- | 3.3 | -- | 4.3 | -- | -- | -- | -- |

*All shown interactions are in Angstroms.*
*† Chain A from PDB 3ZFZ (Otero et al., 2013) with the antibiotic ceftaroline bound to the allosteric and catalytic sites.*
*# Chain A from PDB 3ZG5 (Otero et al., 2013) with the peptidoglycan substrate bound to the allosteric site.*
** All the unique salt bridges described by Otero et al. to connect the allosteric and the catalytic sites are shown with asterisk.*

**Table S5.** Salt bridge interaction networks observed across the room-temperature structures of PBP2a reported in this study, as well as the apo structure (PDB 1VQQ) (Lim & Strynadka, 2002) and the PBP2a structures complexed with the antibiotic ceftaroline (PDB 3ZFZ) (Otero *et al.*, 2013) and peptidoglycan (PDB 3ZG5) (Otero *et al.*, 2013).

| | | PBP2a-LCLS (This study) | | PBP2a-EuXFEL (This study) | | apo-PBP2a (PDB 1VQQ) | | PBP2a-CEF (PDB 3ZFZ) | PBP2a-PG (PDB 3ZG5) |
|---|---|---|---|---|---|---|---|---|---|
| **Residue** | **Residue** | **Chain A** | **Chain B** | **Chain A** | **Chain B** | **Chain A** | **Chain B** | **Chain A†** | **Chain A#** |
| Asp27 | Lys93 | Y | Y | Y | Y | Y | N | Y | Y |
| Asp27 | Lys87 | Y | N | N | N | N | N | N | N |
| Asp35 | Arg83 | Y | Y | Y | Y | Y | Y | Y | Y |
| Glu38 | Arg83 | Y | Y | Y | Y | Y | Y | Y | Y |
| Lys43 | Glu59 | N | Y | Y | Y | Y | Y | Y | Y |
| Asp48 | Lys124 | N | N | N | N | N | Y | N | Y |
| Asp56 | Lys138 | N | N | N | N | N | N | N | Y |
| Lys43 | Glu64 | N | N | N | N | Y | Y | N | Y |
| Glu61 | Arg65 | Y | Y | Y | Y | Y | Y | Y | Y |
| Arg65 | Asp304 | N | Y | Y | Y | N | N | N | N |
| Asp82 | Lys84 | N | N | Y | N | N | Y | N | N |
| Asp82 | Lys100 | N | N | N | N | Y | N | Y | N |
| Lys86 | Asp96 | Y | N | Y | N | Y | N | Y | Y |
| Lys92 | Asp126 | N | N | N | N | N | N | N | Y |
| Arg94 | Asp96 | Y | Y | Y | Y | Y | Y | Y | Y |
| Lys102 | Asp77 | N | N | N | Y | N | N | Y | N |
| Lys100 | Asp109 | Y | N | N | Y | Y | Y | Y | Y |
| Lys124 | Glu119 | N | N | Y | N | N | N | Y | N |
| Lys124 | Asp120 | N | N | Y | N | Y | N | Y | N |
| Lys138 | Asp128 | N | N | N | N | N | N | Y | N |
| Lys148 | Asp295 | Y | Y | Y | Y | Y | Y | Y | Y |
| Arg151 | Asp288 | Y | Y | Y | Y | Y | Y | Y | Y |
| Arg151 | Glu284 | Y | Y | Y | Y | Y | Y | Y | Y |
| Lys153 | Glu150 | N | N | N | N | N | N | Y | N |
| Lys153 | Asp323 | Y | Y | Y | Y | Y | N | Y | N |
| Lys153 | Glu161 | Y | Y | Y | Y | Y | Y | Y | Y |
| Arg157 | Glu356 | Y | Y | Y | Y | Y | Y | Y | Y |
| Arg157 | Asp667 | Y | Y | Y | Y | Y | Y | Y | Y |
| Lys176 | Asp208 | Y | Y | N | Y | N | Y | N | N |
| Lys180 | Glu194 | N | N | N | N | N | N | Y | N |
| Lys181 | Asp182 | N | N | N | N | N | N | N | Y |

| | | | | | | | | | |
|---|---|---|---|---|---|---|---|---|---|
| Lys184 | Glu194 | N | N | N | N | N | N | N | Y |
| Glu194 | Lys198 | N | Y | Y | Y | N | N | N | Y |
| Asp195 | Lys198 | Y | Y | Y | N | Y | N | N | Y |
| Lys198 | Asp202 | N | N | N | N | N | N | Y | N |
| Asp226 | Lys230 | N | Y | N | N | N | Y | N | N |
| Lys219 | Asp221 | N | N | N | N | N | N | Y | Y |
| Arg241 | Glu284 | Y | Y | Y | Y | Y | Y | Y | Y |
| Lys247 | Glu246 | N | N | Y | N | N | N | N | N |
| Lys247 | Asp367 | N | N | Y | N | N | N | Y | Y |
| Glu262 | Lys265 | N | N | N | N | N | N | N | Y |
| Glu263 | Lys280 | Y | Y | Y | Y | Y | Y | Y | Y |
| Glu268 | Lys285 | Y | Y | Y | Y | Y | N | N | Y |
| Glu268 | Lys267 | N | N | Y | N | N | N | N | N |
| Lys273 | Asp275 | N | N | N | Y | N | Y | Y | Y |
| Lys273 | Glu294 | Y | Y | Y | Y | Y | N | Y | N |
| Lys273 | Asp295 | N | N | N | N | N | N | N | Y |
| Asp288 | Lys289 | N | N | Y | Y | N | N | N | N |
| Lys290 | Asp552 | N | Y | N | Y | N | Y | N | Y |
| Glu294 | Lys316 | Y | Y | Y | Y | Y | Y | Y | Y |
| Arg298 | Glu315 | Y | Y | Y | Y | Y | Y | Y | Y |
| Glu315 | Lys317 | Y | Y | N | Y | N | Y | N | N |
| Lys318 | Asp320 | Y | Y | Y | Y | Y | Y | Y | Y |
| Lys319 | Glu294 | Y | Y | Y | Y | Y | Y | Y | Y |
| Asp329 | Lys331 | Y | Y | Y | Y | Y | Y | Y | Y |
| Asp343 | Lys639 | Y | Y | Y | Y | Y | N | Y | N |
| Lys382 | Glu378 | N | Y | N | N | N | Y | N | N |
| Lys382 | Glu385 | N | N | N | N | N | Y | N | N |
| Lys382 | Asp386 | Y | N | Y | N | Y | N | Y | N |
| Lys387 | Asp635 | N | N | N | N | N | N | N | Y |
| Glu389 | Lys634 | N | N | N | N | Y | Y | N | Y |
| Asp420 | Lys422 | Y | N | N | N | N | N | N | N |
| Lys422 | Asp421 | Y | Y | Y | Y | Y | Y | N | N |
| Lys426 | Asp428 | N | N | N | N | Y | N | Y | N |
| Lys434 | Asp435 | Y | Y | Y | Y | N | Y | N | Y |
| Lys434 | Glu511 | Y | Y | Y | Y | Y | Y | Y | Y |

| | | | | | | | | | |
|---|---|---|---|---|---|---|---|---|---|
| Asp435 | Lys436 | N | N | Y | N | N | N | N | Y |
| Arg445 | Asp463 | Y | Y | Y | Y | Y | Y | Y | Y |
| Lys456 | Asp573 | Y | Y | Y | Y | N | Y | N | Y |
| Glu480 | Lys484 | Y | Y | Y | Y | Y | Y | N | N |
| Lys506 | Asp509 | N | N | Y | N | N | N | N | N |
| Lys506 | Glu523 | N | N | Y | N | N | N | N | N |
| Glu523 | Lys604 | N | Y | N | Y | N | Y | N | N |
| Lys559 | Glu490 | Y | Y | Y | Y | Y | Y | Y | Y |
| Glu539 | Lys622 | Y | Y | Y | Y | N | N | N | N |
| Lys565 | Glu566 | Y | N | Y | N | N | N | N | N |
| Lys581 | Glu460 | Y | N | Y | Y | Y | N | Y | Y |
| Lys584 | Glu585 | N | N | N | N | Y | N | N | N |
| Asp586 | Lys647 | Y | Y | Y | Y | Y | Y | N | N |
| Arg589 | Asp654 | Y | Y | Y | Y | Y | Y | Y | Y |
| Arg612 | Asp635 | Y | Y | Y | Y | N | Y | N | Y |
| Arg612 | Glu602 | N | N | N | N | Y | N | Y | N |
| Asp621 | Lys663 | Y | N | Y | N | N | N | N | Y |
| Lys622 | Asp623 | N | N | N | N | N | N | Y | N |
| Asp623 | Lys663 | N | N | Y | N | N | N | N | N |
| Lys634 | Asp635 | N | N | N | N | N | N | Y | N |
| Lys662 | Glu658 | N | Y | Y | Y | Y | Y | Y | Y |
| Lys662 | Glu668 | Y | Y | Y | Y | Y | Y | Y | Y |
| Glu668 | Lys331 | Y | Y | Y | Y | Y | N | N | Y |

*All shown interactions are in Angstroms.*
† *Chain A from PDB 3ZFZ* (Otero *et al.*, 2013) *with the antibiotic ceftaroline bound to the allosteric and catalytic sites.*
### *Chain A from PDB 3ZG5* (Otero *et al.*, 2013) *with the peptidoglycan substrate bound to the allosteric site.*

**Table S6.** Comparison of the salt bridge interactions observed in the room-temperature structures at XFELs and those from the cryogenic structures previously published. All these salt bridges have been extracted from Table S5. The color codes of these interactions match those shown in Figure 7.

| | |
|---|---|
| List of all 18 conserved salt bridge interactions across all structures. | **Asp35-Arg83**<br>**Glu38-Arg83**<br>**Glu61-Arg65**<br>**Arg94-Asp96**<br>**Lys148-Asp295**<br>**Arg151-Asp288**<br>**Arg157-Glu356**<br>**Arg241-Glu284**<br>**Glu263-Lys280**<br>**Glu294-Lys316**<br>**Lys318-Asp320**<br>**Lys319-Glu294**<br>**Lys434-Glu511**<br>**Arg445-Asp463**<br>**Lys559-Glu490**<br>**Asp586-Lys647**<br>**Arg589-Asp654**<br>**Lys622-Glu668** |
| List of 17 novel salt bridges found in either chain of the XFEL structures but not in any of the chains of the cryogenic structure of the apo protein (PDB 1VQQ) (light orange) and the 12 interactions not even described by MD simulations are shown (orange). | **Asp27-Lys87**<br>**Arg65-Asp304**<br>**Lys102-Asp77**<br>**Lys124-Glu119**<br>**Glu194-Lys198**<br>**Lys247-Asp246**<br>**Lys247-Asp367**<br>**Lys267-Glu268**<br>**Asp288-Lys289**<br>**Asp420-Lys422**<br>**Asp435-Lys436**<br>**Lys506-Glu509**<br>**Lys506-Glu523**<br>**Lys539-Lys622**<br>**Lys565-Glu566**<br>**Asp621-Lys663**<br>**Asp623-Lys663** |
| List of 24 salt bridges found in either chain A of XFEL structures but not in the complex of PBP2a with ceftaroline (PDB 3ZFZ) (Otero *et al.*, 2013). These are salt bridges that are broken upon binding. | **Asp27-Lys87**<br>**Asp27-Lys93**<br>**Arg65-Asp304**<br>**Asp48-Lys124**<br>**Lys176-Asp208**<br>**Asp195-Lys198**<br>**Glu194-Lys198**<br>**Lys198-Asp202**<br>**Glu268-Lys285**<br>**Glu268-Lys267**<br>**Asp288-Lys289**<br>**Glu315-Lys317**<br>**Asp420-Lys422**<br>**Lys422-Asp421**<br>**Lys434-Asp435**<br>**Lys456-Asp573**<br>**Glu480-Lys484** |

| | |
|---|---|
| | Lys565-Glu566<br>Asp586-Lys647<br>Arg612-Glu635<br>Asp621-Lys663<br>Glu668-Lys331 |
| List of 12 salt bridges not found in any of the chains A of XFEL structures but seen in chain A of the complex of PBP2a with ceftaroline (PDB 3ZFZ) (Otero *et al.*, 2013). These are salt bridges that are newly formed upon binding. | Asp82-Lys100<br>Lys102-Asp77<br>Lys138-Asp128<br>Lys153-Glu150<br>Lys180-Glu194<br>Lys198-Asp202<br>Lys219-Asp221<br>Lys273-Asp275<br>Lys426-Asp428<br>Arg612-Glu602<br>Lys622-Asp623<br>Lys634-Asp635 |
| List of 22 salt bridges found in either chain A of XFEL structures but not in the complex of PBP2a with PG (PDB 3ZG5) (Otero *et al.*, 2013). These are salt bridges that are broken upon binding. | Asp27-Lys87<br>Arg65-Asp304<br>Asp82-Lys84<br>Lys124-Glu119<br>Lys124-Asp120<br>Lys153-Asp323<br>Lys176-Asp208<br>Lys247-Glu246<br>Glu268-Lys267<br>Lys273-Glu294<br>Asp288-Lys289<br>Glu315-Lys317<br>Asp343-Lys639<br>Lys382-Asp386<br>Lys422-Asp421<br>Glu480-Lys484<br>Lys506-Asp509<br>Lys506-Glu523<br>Glu539-Lys622<br>Lys565-Glu566<br>Asp586-Lys647<br>Asp623-Lys663 |
| List of 13 salt bridges not found in any of the chains A of XFEL structures but seen in chain A of the complex of PBP2a with PG (PDB 3ZG5) (Otero *et al.*, 2013). These are salt bridges that are newly formed upon binding. | Asp48-Lys124<br>Asp56-Lys138<br>Lys43-Glu64<br>Lys92-Asp126<br>Lys181-Asp182<br>Lys184-Glu194<br>Lys219-Asp221<br>Glu262-Lys265<br>Lys273-Asp275<br>Lys273-Asp295<br>Lys290-Asp552<br>Lys387-Asp635<br>Glu389-Lys634 |
| List of all residues involved in the interactions found at the back of PBP2a | Lys290-Asp552<br>Arg157-Glu356<br>Arg157-Asp667 |

| | | |
|---|---|---|
| | | Asp329-Lys331 |
| | | Asp343-Lys639 |
| | | Asp420-Lys422 |
| | | Lys422-Asp421 |
| | | Lys434-Asp435 |
| | | Lys434-Glu511 |
| | | Arg445-Asp463 |
| | | Lys456-Asp573 |
| | | Glu480-Lys484 |
| | | Lys506-Asp509 |
| | | Lys506-Glu523 |
| | | Glu523-Lys604 |
| | | Lys559-Glu490 |
| | | Glu539-Lys622 |
| | | Lys565-Glu566 |
| | | Lys581-Glu460 |
| | | Asp586-Lys647 |
| | | Arg589-Asp654 |
| | | Arg612-Asp635 |
| | | Lys622-Asp623 |
| | | Asp623-Lys663 |
| | | Lys662-Glu658 |
| | | Lys662-Glu668 |
| | | Glu668-Lys331 |
